## Supplementary material for "Repeated loss of variation in insect ovary morphology highlights the role of developmental constraint in life-history evolution": Bibliography of references for ovariole number dataset

### Ovariole number bibliography

1 Department of Organismic and Evolutionary Biology, Harvard University, Cambridge, MA 02138, USA
2 Smithsonian Tropical Research Institute, Panama City, Panama
3 Department of Molecular Genetics and Cell Biology, University of Chicago, Chicago, IL 60637, USA
4 Department of Biology, Universidad de Puerto Rico en Cayey, Cayey 00736, PR
5 Department of Molecular & Cellular Biology, Harvard University, Cambridge, MA 02138, USA

Baker, G. "The biology of A species of Doleschalla (Diptera: Tachinidae), A parasite of Pantorhytes". *Pacific Insects* 19.1-2 (1978): 53–64.

Barbosa, P. and E. Frongillo Jr. "Photoperiod and temperature influences on egg number in *Brachymeria intermedia* (Hymenoptera: Chalcididae), a pupal parasitoid of *Lymantria dispar* (Lepidoptera: Lymantriidae)". *Journal of The New York Entomological Society* 87.2 (1979): 175–180.

Barratt, B., A. Evans, D. Stoltz, S. Vinson, and R. Easingwood. "Virus-like Particles in the Ovaries of *Microctonus aethiopoides* Loan (Hymenoptera: Braconidae), a Parasitoid of Adult Weevils (Coleoptera: Curculionidae)". *Journal of Invertebrate Pathology* 73.2 (1999): 182–188.

Bellinger, R. G. and R. L. Pienkowski. "Interspecific variation in ovariole number in Melanopline grasshoppers (Orthoptera: Acrididae)". *Annals of The Entomological Society of America* 78.1 (1985): 127–130.

Bellinger, R. G. and R. L. Pienkowski. "Non-random resorption of oocytes in grasshoppers (Orthoptera: Acrididae)". *The Canadian Entomologist* 117.9 (1985): 1067–1069.

Benham Jr, G. S. "Digestive and reproductive systems of *Eriborus molestae* Uchida (Hymenoptera: Ichneumonidae)". *International Journal of Insect Morphology and Embryology* 1.2 (1972): 153–161.

Berkebile, D. R., A. P. Weinhold, and D. B. Taylor. "A new method for collecting clean stable fly (Diptera: Muscidae) pupae of known age". *Southwestern Entomologist* 34.4 (2009): 469–477.

Biani, N. B. and W. T. Wcislo. "Notes on the reproductive morphology of the parasitic bee *Megalopta byroni* (Hymenoptera: Halictidae), and a tentative new host record". *Journal of The Kansas Entomological Society* 80.4 (2007): 392–395.

Biliński, S. and B. Petryszak. "The ultrastructure and function of follicle cells in *Foucartia squamulata* (Herbst) (Curculionidae)". *Cell and Tissue Research* 189.2 (1978): 347–353.

Biliński, S. "Oogenesis in *Acerentomon gallicum* Jonescu (Protura)". *Cell and Tissue Research* 179.3 (1977): 401–412.

Biliński, S. "Oogenesis in *Campodea* sp. (Diplura)". *Cell and Tissue Research* 202.1 (1979): 133–143.

Bilinski, S. M. and J. Büning. "Structure of ovaries and oogenesis in the snow scorpionfly *Boreus hyemalis* (Linne) (Mecoptera: Boreidae)". *International Journal of Insect Morphology and Embryology* 27.4 (1998): 333–340.

Biliński, S. M. and T. Szklarzewicz. "The ovary of *Catajapyx aquilonaris* (Insecta, Entognatha): ultrastructure of germarium and terminal filament". *Zoomorphology* 112.4 (1992): 247–251.

Blowers, V., et al. "Notes on the female reproductive system of the South African citrus psylla, *Trioza erytreae* (Del Guercio) (Homoptera: Psyllidae)". *Journal of The Entomological Society of Southern Africa* 30.1 (1967): 75–81.

Bong, L.-J., K.-B. Neoh, C.-Y. Lee, and Z. Jaal. "Effect of diet quality on survival and reproduction of adult *Paederus fuscipes* (Coleoptera: Staphylinidae)". *Journal of Medical Entomology* 51.4 (2014): 752–759.

Bourchier, R. "Growth and development of *Compsilura concinnata* (Meigan) (Diptera: Tachinidae) parasitizing gypsy moth larvae feeding on tannin diets". *The Canadian Entomologist* 123.5 (1991): 1047–1055.

Braman, S. and K. Yeargan. "Reproductive strategies of primary parasitoids of the green cloverworm (Lepidoptera: Noctuidae)". *Environmental Entomology* 20.1 (1991): 349–353.

Brandmayr, P. and T. Zetto-Brandmayr. "The evolution of parental care phenomena in Pterostichine ground beetles, with special reference to the genera *Abax* and *Molops* (Coleoptera, Carabidae)". *Miscellaneous Papers Landbouwhogeschool. Wageningen* 18 (1979): 35–49.

Branson, D. H. "Influence of individual body size on reproductive traits in melanopline grasshoppers (Orthoptera: Acrididae)". *Journal of Orthoptera Research* 17.2 (2008): 259–264.

Branson, D. H. "Life-history responses of *Ageneotettix deorum* (Scudder) (Orthoptera: Acrididae) to host plant availability and population density". *Journal of The Kansas Entomological Society* 79.2 (2006): 146–156.

Brent, C. S. and J. F. Traniello. "Effect of enhanced dietary nitrogen on reproductive maturation of the termite *Zootermopsis angusticollis* (Isoptera: Termopsidae)". *Environmental Entomology* 31.2 (2002): 313–318.

Brent, C. S., J. F. Traniello, and E. L. Vargo. "Benefits and costs of secondary polygyny in the dampwood termite *Zootermopsis angusticollis*". *Environmental Entomology* 37.4 (2008): 883–888.

Brothers, D. J. "Alternative life-history styles of mutillid wasps (Insecta, Hymenoptera)". *Alternative life-history styles of animals* (1989): 279–291.

Browne, J. and C. H. Scholtz. “Evolution of the scarab hindwing articulation and wing base: a contribution toward the phylogeny of the Scarabaeidae (Scarabaeoidea: Coleoptera)”. *Systematic Entomology* 23.4 (1998): 307–326.

Brunt, A. M. “The histology of the first batch of eggs and their associated tissues in the ovariole of *Dysdercus fasciatus* Signoret (Heteroptera: Pyrrhocoridae) as seen with the light microscope”. *Journal of Morphology* 134.1 (1971): 105–129.

Bugnion, E. “Le *Cissites testaceus* Vab. des Indes et de Ceylon”. *Bulletin de la Société entomologique d’Égypte* 1.8 (1909): 182–204.

Bugnion, É. and N. Popoff. “Anatomie de la reine et du roi-termite”. *Mémoires de La Société Zoologique de France* 25 (1912): 210–232.

Büning, J. and S. Sohst. “Ultrastructure and cluster formation in ovaries of bark lice, *Peripsocus phaeopterus* (Stephens) and *Stenopsocus stigmaticus* (Imhof and Labram) (Insecta: Psocoptera)”. *International Journal of Insect Morphology and Embryology* 19.5-6 (1990): 227–241.

Büning, J. “Morphology, ultrastructure, and germ cell cluster formation in ovarioles of aphids”. *Journal of Morphology* 186.2 (1985): 209–221.

Büning, J. “Ovariole structure supports sistergroup relationship of Neuropterida and Coleoptera”. *Arthropod Systematics and Phylogeny* 64.2 (2006): 115–126.

Büning, J. “The ovary of *Raphidia flavipes* is telotrophic and of the *Sialis* type (Insecta, Raphidioptera)”. *Zoomorphologie* 95.2 (1980): 127–131.

Büning, J. “The telotrophic nature of ovarioles of polyphage Coleoptera”. *Zoomorphologie* 93.1 (1979): 51–57.

Büning, J. “The telotrophic ovary known from Neuropterida exists also in the myxophagan beetle *Hydroscapha natans*”. *Development Genes and Evolution* 215.12 (2005): 597–607.

Büning, J. “The trophic tissue of telotrophic ovarioles in polyphage Coleoptera”. *Zoomorphologie* 93.1 (1979): 33–50.

Büning, J. and S. Sohst. “The flea ovary: ultrastructure and analysis of cell clusters”. *Tissue and Cell* 20.5 (1988): 783–795.

Butler, L. “Biology of the half-wing geometer, *Phigalia titea* Cramer (Geometridae), as a member of a looper complex in West Virginia”. *Jounral Lepidopterists Society* 39 (1985): 177–186.

Calder, A. A. “Gross morphology of the soft parts of the male and female reproductive systems of Curculionoidea (Coleoptera)”. *Journal of Natural History* 24.2 (1990): 453–505.

Callahan, P. S. “Serial morphology as a technique for determination of reproductive patterns in the corn earworm, *Heliothis zea* (Boddie)”. *Annals of The Entomological Society of America* 51.5 (1958): 413–428.

Cao, S., S. Shang, Y. Zhang, et al. “Anatomy of the reproductive system of *Pieris melete* Ménétriès (Lepidoptera: Pieridae).” *Journal of Northwest A and F University-Natural Science Edition* 40.9 (2012): 77–82.

Capman, W. C. “Natural history of the common sooty wing skipper, *Pholisora catullus* (Lepidoptera: Hesperiidae), in Central Illinois”. *The Great Lakes Entomologist* 23.3 (2017): 151–157.

Carl, K. “The natural enemies of the pear-slug, *Caliroa cerasi* (L.) (Hym., Tenthredinidae), in Europe”. *Zeitschrift Für Angewandte Entomologie* 80.1-4 (1976): 138–161.

Carlberg, U. “Evolutionary and ecological aspects on ovarian diversity in Phasmida (Insecta).” *Zoologische Jahrbücher. Abteilung für Systematic Okologie und Geographie der Tiere. Jena* 114.1 (1987): 45–63.

Carlberg, U. “Ovary anatomy in Phasmida (Insecta)”. *Zoologische Jahrbücher. Abteilung für Systematic Okologie und Geographie der Tiere. Jena* 115.1 (1987): 77–84.

Carlberg, U. “Ovary anatomy in Phasmida (Insecta). II”. *Zoologische Jahrbücher. Abteilung für Systematic Okologie und Geographie der Tiere. Jena* 118.1 (1989): 9–14.

Cassidy, J. D. and R. C. King. “Ovarian development in *Habrobracon juglandis* (Ashmead) (Hymenoptera: Braconidae). I. The origin and differentiation of the oocyte nurse cell complex”. *The Biological Bulletin* 143.3 (1972): 483–505.

Catts, E. “Field Behavior of Adult *Cephenemyia* (Diptera: Oestridae) 1, 2”. *The Canadian Entomologist* 96.3 (1964): 579–585.

Cave, M. D. “Absence of amplification of ribosomal DNA in the polytrophic meroistic ovary of the giant silkworm moth, *Antheraea pernyi* (Lepidoptera: Saturniidae)”. *Wilhelm Roux's Archives of Developmental Biology* 184.2 (1978): 135–142.

Cervera, A., A. C. Maymo, R. Martínez-Pardo, and M. D. Garcerá. “Vitellogenesis inhibition in *Oncopeltus fasciatus* females (Heteroptera: Lygaeidae) exposed to cadmium”. *Journal of Insect Physiology* 51.8 (2005): 895–911.

Chanda, S. and S. Chakravorty. “Morphogenetic derangements in the reproductive system of *Bracon hebetor*, a beneficial parasitoid, bred on juvenoid treated host (Corcyra cephalonica) larvae”. *Indian Journal of Experimental Biology* 38 (2000): 700–794.

Charlwood, J. d. and J. A. Rafael. “Autogeny in the river negro horse fly, *Lepiselaga crassipes*, and un undescribed species of *Stenotabanus* (Diptera: Tabanidae) from Amazonas, Brazil”. *Journal of Medical Entomology* 17.6 (1980): 519–521.

Chiang, R. “Functional anatomy of the vagina muscles in the adult western conifer seed bug, *Leptoglossus occidentalis* (Heteroptera: Coreidae), and its implication for the egg laying behaviour in insects”. *Arthropod Structure and Development* 39.4 (2010): 261–267.

Chiu, M.-C., C.-G. Huang, W.-J. Wu, and S.-F. Shiao. “Morphological allometry and intersexuality in horsehair-worm-infected mantids, *Hierodula formosana* (Mantodea: Mantidae)”. *Parasitology* 142.8 (2015): 1130–1142.

Clift, A. and F. McDonald. “Morphology of the internal reproductive system of *Lucilia cuprina* (Wied.) (Diptera: Calliphoridae) and a method of determining the age of both sexes”. *International Journal of Insect Morphology and Embryology* 2.4 (1973): 327–333.

Coninck, E. de and R. Coessens. “The structure of the internal genitalia of *Acrotrichis intermedia* (Gillm., 1845) (Col. Ptiliidae)”. *Deutsche Entomologische Zeitschrift* 29.1-3 (1982): 51–55.

Coombs, M. T. “Influence of adult food deprivation and body size on fecundity and longevity of *Trichopoda giacomellii*: A South American parasitoid of *Nezara viridula*”. *Biological Control* 8.2 (1997): 119–123.

Cooper, K. W. “A southern Californian *Boreus*, *B. notoperates* n. sp. I. *Comparative morphology* and systematics (Mecoptera: Boreidae)”. *Psyche: A Journal of Entomology* 79.4 (1972): 269–283.

Cooper, K. W. “The genital anatomy and mating behavior of *Boreus brumalis* Fitch (Mecoptera)”. *American Midland Naturalist* 23.2 (1940): 354–367.

Coppel, H. and M. Maw. “Studies on dipterous parasites of the spruce budworm, *Choristoneura fumiferana* (Clem.) (Lepidoptera: Tortricidae): III. *Ceromasia auricaudata* tns. (Diptera: Tachinidae)”. *Canadian Journal of Zoology* 32.3 (1954): 144–156.

Cox, M. and D. Windsor. “The first instar larva of *Aulacoscelis appendiculata* n. sp. (Coleoptera: Chrysomelidae: Aulacoscelinae) and its value in the placement of the Aulacoscelinae”. *Journal of Natural History* 33.7 (1999): 1049–1087.

Craddock, E. M. and M. P. Kambysellis. “Adaptive radiation in the Hawaiian *Drosophila* (Diptera: Drosophilidae): Ecological and reproductive character analyses”. *Pacific Science* 51.4 (1997): 475–489.

Cruz-Landim, C. d., R. D. Reginato, and V. L. Imperatriz-Fonseca. “Variation on ovariole number in Meliponinae (Hymenoptera, Apidae) queen's ovaries, with comments on ovary development and caste differentiation”. *Papeis Avulsos de Zoologia (São Paulo)* 40 (1998): 289–296.

Danks, H. “Seasonal cycle and biology of *Winthemia rufopicta* (Diptera: Tachinidae) as a parasite of *Heliothis* spp. (Lepidoptera: Noctuidae) on tobacco in North Carolina”. *The Canadian Entomologist* 107.6 (1975): 639–654.

Daumal, J. and H. Boinel. “Variability in fecundity and plasticity of oviposition behavior in *Anagasta kuehniella* (Lepidoptera: Pyralidae)”. *Annals of The Entomological Society of America* 87.2 (1994): 250–256.

Davies, R. “The postembryonic development of the female reproductive system in *Limothrips cerealium* Haliday (Thysanoptera: Thripidae)”. *Proceedings of the Zoological Society of London* 136.3 (1961): 411–437.

Davis, N. T. and R. L. Usinger. “The biology and relationships of the Joppeicidae (Heteroptera)”. *Annals of The Entomological Society of America* 63.2 (1970): 577–587.

Dhileepan, K. and T. Ananthakrishnan. "Ovarian polymorphism in relation to reproductive diversity and associated histological and histochemical attributes in some sporophagous tubuliferan Thysanoptera". *Proceedings: Animal Sciences* 96.1 (1987): 1–13.

Diefenbach, L. M. G., L. R. Redaelli, and D. N. Gassen. "Characterization of the internal reproductive organs and their state as diapause indicator in *Phytalus sanctipauli* Blanchard, 1850 (Coleoptera, Scarabaeidae)". *Revista Brasileira de Biologia* 58.3 (1998): 541–546.

Dinan, L. "Ecdysteroids in adults and eggs of the house cricket, *Acheta domesticus* (Orthoptera: Gryllidae)". *Comparative Biochemistry and Physiology Part B: Biochemistry and Molecular Biology* 116.2 (1997): 129–135.

Dowell, R. "Ovary structure and reproductive biologies of larval parasitoids of the alfalfa weevil (Coleoptera: Curculionidae) 1 2". *The Canadian Entomologist* 110.5 (1978): 507–512.

Dowell, R. V. and D. J. Horn. "Adaptive strategies of larval parasitoids of the alfalfa weevil (Coleoptera: Curculionidae)". *The Canadian Entomologist* 109.5 (1977): 641–648.

Dowell, R. V. and T. K. Wood. "Ovary structure, fecundity and mating receptivity in *Umbonia crassicornis* Amyot and Serville 1843 (Hemiptera: Membracidae)". *The Pan-Pacific Entomologist* 85.4 (2009): 187–190.

Dunham, R. S. "A life history study of *Caecilius aurantiacus* (Hagen) (Psocoptera: Caeciliidae)". *The Great Lakes Entomologist* 5.1 (2017): 17–27.

Dunlap-Pianka, H., C. L. Boggs, and L. E. Gilbert. "Ovarian dynamics in heliconiine butterflies: programmed senescence versus eternal youth". *Science* 197.4302 (1977): 487–490.

Dutrillaux, A. M., D. Pluot-Sigwalt, and B. Dutrillaux. "Ovo-viviparity in the darkling beetle, *Alegoria castelnaui* (Tenebrioninae: Ulomini), from Guadeloupe." *European Journal of Entomology* 107.4 (2010): 481–485.

Edmonds, W. "Internal anatomy of *Coprophanaeus lancifer* (L.) (Coleoptera: Scarabaeidae)". *International Journal of Insect Morphology and Embryology* 3.2 (1974): 257–272.

Eguagie, W. E. "*PhD Thesis*: Studies on the ecology of thistle lace bugs, in particular *Tingis ampliata* (Heteroptera: Tingidae)". Diss. Imperial College London, 1969.

Eidmann, H. "Morphologische und physiologische Untersuchungen am weiblichen Genitalapparat der Lepidopteren: I. Morphologischer Teil". *Zeitschrift Für Angewandte Entomologie* 15.1 (1929): 1–66.

Elamin, A. E., A. M. Abdalla, A. M. El Naim, et al. "The biology of senegalese grasshopper (*Oedaleus senegalensis*, Krauss, 1877) (Orthoptera: Acrididae)". *International Journal of Advances in Life Science and Technology* 1.1 (2014): 6–15.

Elelimy, H., N. Ghazawy, A. Omar, and A. Meguid. "Morphology, histology and ovary development of the female reproductive system of *Spilostethus pandurus* (Scopoli) (Hemiptera: Lygaeidae)". *African Entomology* 25.2 (2017): 515–523.

Eliopoulos, P. A., J. A. Harvey, C. G. Athanassiou, and G. J. Stathas. "Effect of biotic and abiotic factors on reproductive parameters of the synovigenic endoparasitoid *Venturia canescens*". *Physiological Entomology* 28.4 (2003): 268–275.

Elliott, H. and R. Bashford. "The life history of *Mnesampela privata* (Guen.) (Lepidoptera: Geometridae) a defoliator of young eucalypts". *Australian Journal of Entomology* 17.3 (1978): 201–204.

Emeljanov, A., N. Golub, and V. Kuznetsova. "Birdlice, and Sucking Lice (Psocoptera, Phthiraptera: Mallophaga, Anoplura)". *Entomological Review* 81.7 (2001): 767–785.

Etman, A. A. and G. Hooper. "Developmental and reproductive biology of *Spodoptera litura* (F.) (Lepidoptera: Noctuidae)". *Australian Journal of Entomology* 18.4 (1980): 363–372.

Faille, A. and D. Pluot-Sigwalt. "Convergent reduction of ovariole number associated with subterranean life in beetles". *PloS One* 10.7 (2015): 1–14.

Farder-Gomes, C. F., H. C. P. Santos, M. A. Oliveira, J. C. Zanuncio, and J. E. Serrão. "Morphology of ovary and spermathecae of the parasitoid *Eibesfeldtphora tonhascai* Brown (Diptera: Phoridae)". *Protoplasma* 256.1 (2019): 3–11.

Farrow, R. “Population dynamics of the Australian plague locust, *Chortoicetes terminifera* (Walker) in central western New South Wales. II. Factors influencing natality and survival.” *Australian Journal of Zoology* 30.2 (1982): 199–222.

Fátima Ribeiro, M. de, P. de Souza Santos-Filho, and V. L. Imperatriz-Fonseca. “Size variation and egg laying performance in *Plebeia remota* queens (Hymenoptera, Apidae, Meliponini)”. *Apidologie* 37.6 (2006): 653–664.

Fatzinger, C. W. “Morphology of the reproductive organs of *Dioryctria abietella* (Lepidoptera: Pyralidae (Phycitinae))”. *Annals of The Entomological Society of America* 63.5 (1970): 1256–1261.

Ferrar, P. “Macrolarviparous reproduction in Euphumosia (Diptera: Calliphoridae)”. *Australian Journal of Entomology* 17.1 (1978): 13–17.

Fink, T. J., T. Soldán, J. G. Peters, and W. L. Peters. “The reproductive life history of the predacious, sand-burrowing mayfly *Dolania americana* (Ephemeroptera: Behningiidae) and comparisons with other mayflies”. *Canadian Journal of Zoology* 69.4 (1991): 1083–1093.

Fisher, R. M. and B. J. Sampson. “Morphological specializations of the bumble bee social parasite *Psithyrus ashtoni* (Cresson) (Hymenoptera: Apidae)”. *The Canadian Entomologist* 124.1 (1992): 69–77.

Fitt, G. P. “Comparative fecundity, clutch size, ovariole number and egg size of *Dacus tryoni* and *D. jarvisi*, and their relationship to body size”. *Entomologia Experimentalis Et Applicata* 55.1 (1990): 11–21.

Fitt, G. P. “Variation in ovariole number and egg size of species of *Dacus* (Diptera; Tephritidae) and their relation to host specialization”. *Ecological Entomology* 15.3 (1990): 255–264.

Flerrar, P. “Macrolarviparous reproduction in Ameniinae (Diptera: Calliphoridae)”. *Systematic Entomology* 1.2 (1976): 107–116.

Force, D. C. and M. L. Thompson. “Parasitoids of the immature stages of several southwestern yucca moths”. *The Southwestern Naturalist* 29.1 (1984): 45–56.

Fortes, P., G. Salvador, and F. Consoli. “Ovary development and maturation in *Nezara viridula* (L.) (Hemiptera: Pentatomidae)”. *Neotropical Entomology* 40.1 (2011): 89–96.

Freeman, B. and A. Geoghgen. “Size and fecundity in the Jamaican gall-midge *Asphondylia boerhaaviae*”. *Ecological Entomology* 12.3 (1987): 239–249.

Fremlin, M. “Single mothers: Minotaur beetle females *Typhaeus typhoeus* (L.) (Coleoptera: Geotrupidae) nest on their own”. *Bulletin of the Amateur Entomologists' Society* 76 (2017): 87–95.

Gaikwad, S. M., Y. J. Koli, and G. P. Bhawane. “Histomorphology of the Female Reproductive System in *Papilio polytes polytes* Linnaeus, 1758 (Lepidoptera: Papilionidae)”. *Proceedings of The National Academy of Sciences, India Section B: Biological Sciences* 84.4 (2014): 901–908.

Gaines, D. N. “*PhD Thesis*: Studies on *Conura torvina* (Hymenoptera: Chalcididae) reproduction and biology in relation to hosts in Brassica Crops.” Diss. Virginia Tech, 1997.

Gaino, E. and M. Rebora. “Detection of apoptosis in the ovarian follicle cells of *Ecdyonurus venosus* (Ephemeroptera, Heptageniidae)”. *Italian Journal of Zoology* 70.4 (2003): 291–295.

Galbraith, D. and C. Fernando. “The life history of *Gerris remigis* (Heteroptera: Gerridae) in a small stream in southern Ontario”. *The Canadian Entomologist* 109.2 (1977): 221–228.

Garbiec, A., J. Kubrakiewicz, M. Mazurkiewicz-Kania, B. Simiczyjew, and I. Jkedrzejowska. “Asymmetry in structure of the eggshell in *Osmylus fulvicephalus* (Neuroptera: Osmylidae): an exceptional case of breaking symmetry during neuropteran oogenesis”. *Protoplasma* 253.4 (2016): 1033–1042.

Gardner, J. A. “Revision of the genera of the tribe Stigmoderini (Coleoptera: Buprestidae) with a discussion of phylogenetic relationships”. *Invertebrate Systematics* 3.3 (1989): 291–361.

Gauld, I. D., D. B. Wahl, and G. R. Broad. “The suprageneric groups of the Pimplinae (Hymenoptera: Ichneumonidae): a cladistic re-evaluation and evolutionary biological study”. *Zoological Journal of The Linnean Society* 136.3 (2002): 421–485.

Genc, H. and J. L. Nation. “An artificial diet for the butterfly *Phyciodes phaon* (Lepidoptera: Nymphalidae)”. *Florida Entomologist* 87.2 (2004): 194–199.

Gerber, G. "Reproductive behaviour and physiology of *Tenebrio molitor* (Coleoptera: Tenebrionidae): II. Egg development and oviposition in young females and the effects of mating". *The Canadian Entomologist* 107.5 (1975): 551–559.

Gerber, G. and N. Church. "The reproductive cycles of male and female *Lytta nuttalli* (Coleoptera: Meloidae) 1 3". *The Canadian Entomologist* 108.10 (1976): 1125–1136.

Gerber, G., N. Church, and J. Rempel. "The anatomy, histology, and physiology of the reproductive systems of *Lytta nuttalli* Say (Coleoptera: Meloidae). I. The internal genitalia". *Canadian Journal of Zoology* 49.4 (1971): 523–533.

Gerber, G., G. Neill, and P. Westdal. "The anatomy and histology of the internal reproductive organs of the sunflower beetle, *Zygogramma exclamationis* (Coleoptera: Chrysomelidae)". *Canadian Journal of Zoology* 56.12 (1978): 2542–2553.

Gerling, D. and H. Hermann. "The oviposition and life cycle of *Anthrax tigrinus*,[Dipt.: Bombyliidae] a parasite of carpenter bees [Hym.: Xylocopidae]". *Entomophaga* 21.3 (1976): 227–233.

Ghoneim, K. S. "Embryonic and postembryonic development of blister beetles (Coleoptera: Meloidae) in the world: A synopsis". *International Journal of Biological Sciences* 2 (2013): 6–18.

Gil-Fernandez, C. and L. Black. "Some aspects of the internal anatomy of the leafhopper *Agallia constricta* (Homoptera: Cicadellidae)". *Annals of The Entomological Society of America* 58.3 (1965): 275–284.

Gnatzy, W., W. Volknandt, and A. Dzwoneck. "Egg-laying behavior and morphological and chemical characterization of egg surface and egg attachment glue of the digger wasp *Ampulex compressa* (Hymenoptera, Ampulicidae)". *Arthropod Structure and Development* 47.1 (2018): 74–81.

Goldson, S., M. McNeill, J. Proffitt, and A. Hower. "An investigation into the reproductive characteristics of *Microctonus hyperodae* (Hym.: Braconidae), a parasitoid of *Listronotus bonariensis* (Col.: Curculionidae)". *Entomophaga* 40.3-4 (1995): 413–426.

Golub, N. and S. Nokkala. "Chromosome numbers of two sucking louse species (Insecta, Phthiraptera, Anoplura)". *Hereditas* 141.1 (2004): 94–96.

Gottanka, J. and J. Büning. "Mayflies (Ephemeroptera), the most "primitive" winged insects, have telotrophic meroistic ovaries". *Roux's Archives of Developmental Biology* 203.1-2 (1993): 18–27.

Gottanka, J. and J. Büning. "Oocytes develop from interconnected cystocytes in the panoistic ovary of *Nemoura* sp. (Pictet) (Plecoptera: Nemouridae)". *International Journal of Insect Morphology and Embryology* 19.5-6 (1990): 219–225.

Gôukon, K., Y. Maeta, and S. F. Sakagami. "Seasonal changes in ovarian state in a eusocial halictine bee, *Lasioglossum duplex*, based on stages of the oldest oocytes in each ovariole (Hymenoptera: Halictidae)". *Researches On Population Ecology* 29.2 (1987): 255–269.

Gower, A. "A study of *Limnephilus lunatus* Curtis (Trichoptera: Limnephilidae) with reference to its life cycle in watercress beds". *Transactions of The Royal Entomological Society of London* 119.10 (1967): 283–302.

Grandi, G., R. Barbieri, and G. Colombo. "Oogenesis in *Kalotermes flavicollis* (Fabr.) (Isoptera: Kalotermitidae). I. Differentiation and maturation of oocytes in female supplementary reproductives". *Italian Journal of Zoology* 55.1-4 (1988): 103–122.

Gray, B. and K. Lamb. "Some observations on the Malpighian tubules and ovarioles in *Myrmecia dispar* (Clark) (Hymenoptera: Formicidae)". *Australian Journal of Entomology* 7.1 (1968): 80–81.

Grenier, A.-M. and P. Nardon. "The genetic control of ovariole number in *Sitophilus oryzae* L (Coleoptera, Curculionidae) is temperature sensitive". *Genetics Selection Evolution* 26.5 (1994): 1–18.

Griffith, C. and J. Lai-Fook. "The ovaries and changes in their structural components at the end of vitellogenesis and during vitelline membrane formation in the butterfly, Calpodes". *Tissue and Cell* 18.4 (1986): 575–588.

Grodowitz, M. J., T. D. Center, and J. E. Freedman. "A physiological age-grading system for *Neochetina eichhorniae* (Warner) (Coleoptera: Curculionidae), a biological control agent of water hyacinth, *Eichhornia crassipes* (Mart.) Solms." *Biological Control* 9.2 (1997): 89–105.

Grosch, D. S., R. G. Kratsas, and R. M. Petters. "Variation in *Habrobracon juglandis* ovariole number: I. Ovariole number increase induced by extended cold shock of fourth-instar larvae". *Development* 40.1 (1977): 245–251.

Grozeva, S. and N. Simov. "Cytogenetic Studies of Bryocorinae Baerensprung, 1860 True Bugs (Heteroptera: Miridae)". *Acta Zool Bulg Suppl* 2 (2008): 61–70.

Grozeva, S. and V. Kuznetsova. "Karyotypes and some structural properties of the reproductive system of bugs of the subfamily Artheneinae (Heteroptera, Pentatomomorpha, Lygaeidae)". *Entomological Research* 69 (1990): 14–26.

Habluetzel, A., F. Carnevali, L. Lucantoni, L. Grana, A. R. Attili, F. Archilei, M. Antonini, A. Valbonesi, V. Abbadessa, F. Esposito, et al. "Impact of the botanical insecticide Neem Azalon survival and reproduction of the biting louse *Damalinia limbata* on angora goats". *Veterinary Parasitology* 144.3-4 (2007): 328–337.

Hafeez, M. and B. Gardiner. "The internal morphology of the adult of *Tribolium anaphe* Hinton (Coleoptera: Tenebrionidae)". *Proceedings of the Royal Entomological Society of London. Series A, General Entomology* 39.10-12 (1964): 137–145.

Halffter, G., W. D. Edmonds, et al. "The nesting behavior of dung beetles (Scarabaeinae). An ecological and evolutive approach." *Journal of the New York Entomological Society* 91.4 (1982): 512–515.

Halffter, G., C. Huerta, R. de Ma. Ribeiro Sarges, and A. D. Rojas. "Reversal to a Two-Ovaries State in Scarabaeinae (Coleoptera: Scarabaeidae)". *The Coleopterists Bulletin* 67.2 (2013): 94–96.

Hallman, G., C. Morales, and M. Duque. "Biology of *Acrosternum marginatum* (Heteroptera: Pentatomidae) on common beans". *Florida Entomologist* 75.2 (1992): 190–196.

Hatakeyama, M., M. Sawa, and K. Oishi. "Ovarian development and vitellogenesis in the sawfly, *Athalia rosae ruficornis* Jakovlev (Hymenoptera, Tenthredinidae)". *Invertebrate Reproduction and Development* 17.3 (1990): 237–245.

Heather, N. "Studies on female genitalia of Queensland Phasmida". *Australian Journal of Entomology* 4.1 (1965): 33–38.

Hébert, C., C. Cloutier, J. Régnière, and D. F. Perry. "Seasonal biology of *Winthemia fumiferanae* Toth. (Diptera: Tachinidae), a larval–pupal parasitoid of the spruce budworm (Lepidoptera: Tortricidae)". *Canadian Journal of Zoology* 67.10 (1989): 2384–2391.

Heming-van Battum, K. E. and B. Heming. "Structure, function and evolution of the reproductive system in females of *Hebrus pusillus* and *H. ruficeps* (Hemiptera, Gerromorpha, Hebridae)". *Journal of Morphology* 190.2 (1986): 121–167.

Heming, B. S. "History of the germ line in male and female thrips". *Thrips biology and management* 276 (1995): 505–535.

Heming, B. and E. Huebner. "Development of the germ cells and reproductive primordia in male and female embryos of *Rhodnius prolixus* Staal (Hemiptera: Reduviidae)". *Canadian Journal of Zoology* 72.6 (1994): 1100–1119.

Hemptinne, E. B. and Jean-Louis. "Development of ovaries, allometry of reproductive traits and fecundity of *Episyrphus balteatus* (Diptera: Syrphidae)". *European Journal of Entomology* 97 (2000): 165–170.

Henry, C. S. "Eggs and rapagula of *Ululodes* and *Ascaloptynx* (Neuroptera: Ascalaphidae): a comparative study". *Psyche: A Journal of Entomology* 79.1-2 (1972): 1–22.

Henry, E. V., A. Alvarez-Zapata, A. C. Gonzales, E. F. Martins, L. C. Martinez, and J. E. Serrao. "Anatomy and histology of the alimentary canal and ovarioles of *Ceraeochrysa cubana* adults". *Bulletin of Insectology* 70.2 (2017): 181–188.

Hernandez, L. C., G. Fajardo, L. S. Fuentes, and L. Comoglio. "Biology and reproductive traits of *Drymoea veliterna* (Druce, 1885) (Lepidoptera: Geometridae)". *Journal of Insect Biodiversity* 5.12 (2017): 1–9.

Higley, L. G. "Morphology of Reproductive Structures in *Cicindela repanda* (Coleoptera: Cicindelidae)". *Journal of The Kansas Entomological Society* 59.2 (1986): 303–308.

Hill, R. I., C. M. Penz, and P. DeVries. "Phylogenetic analysis and review of *Panacea* and *Batesia* butterflies (Nymphalidae)". *Journal of The Lepidopterists' Society* 56.4 (2002): 199–215.

Hoc, T. and T. Wilkes. “Age determination in the blackfly *Simulium woodi*, a vector of onchocerciasis in Tanzania”. *Medical and Veterinary Entomology* 9.1 (1995): 16–24.

Hodin, J. “Plasticity and constraints in development and evolution”. *Journal of Experimental Zoology* 288.1 (2000): 1–20.

Hodin, J. “She shapes events as they come: plasticity in female insect reproduction”. *Phenotypic plasticity of insects : mechanisms and consequences*. Edited by D. W. Whitman and T. N. Ananthakrishnan. Enfield, NH: Science Publishers, 2009. 423–521.

Hodin, J. and L. M. Riddiford. “Different mechanisms underlie phenotypic plasticity and interspecific variation for a reproductive character in drosophilids (Insecta: Diptera)”. *Evolution* 54.5 (2000): 1638–1653.

Hopkins, J., C. Steelman, and C. Carlton. “Anatomy of the adult female lesser mealworm *Alphitobius diaperinus* (Coleoptera: Tenebrionidae) reproductive system”. *Journal of The Kansas Entomological Society* 65.3 (1992): 299–307.

Houston, T. F. “Observations of the biology and immature stages of the sandgroper *Cylindraustralia kochii* (Saussure), with notes on some congeners (Orthoptera: Cylindrachetidae)”. *Records-Western Australian Museum* 23.3 (2007): 219–234.

Huenefeld, F. and N. P. Kristensen. “The female postabdomen and internal genitalia of the basal moth genus *Agathiphaga* (Insecta: Lepidoptera: Agathiphagidae): morphology and phylogenetic implications”. *Zoological Journal of The Linnean Society* 159.4 (2010): 905–920.

Hummel, N. A., F. G. Zalom, and C. Y. Peng. “Anatomy and histology of reproductive organs of female *Homalodisca coagulata* (Hemiptera: Cicadellidae: Proconiini), with special emphasis on categorization of vitellogenic oocytes”. *Annals of The Entomological Society of America* 99.5 (2006): 920–932.

Hünefeld, F. and N. P. Kristensen. “The female postabdomen and genitalia of the basal moth family Heterobathmiidae (Insecta: Lepidoptera): Structure and phylogenetic significance”. *Arthropod Structure and Development* 41.4 (2012): 395–407.

Ichikawa, Y. and M. Watanabe. “Changes in the number of eggs loaded in *Pantala flavescens* females with age from mass flights (Odonata: Libellulidae)”. *Zoological Science* 31.11 (2014): 721–725.

Ikeda, H., T. Kagaya, K. Kubota, and T. Abe. “Evolutionary relationships among food habit, loss of flight, and reproductive traits: life-history evolution in the Silphinae (Coleoptera: Silphidae)”. *Evolution: International Journal of Organic Evolution* 62.8 (2008): 2065–2079.

Imperatriz-Fonseca, V. L., C. d. Cruz-Landim, and R. S. de Moraes. “Dwarf gynes in *Nannotrigona testaceicornis* (Apidae, Meliponinae, Trigonini). Behaviour, exocrine gland morphology and reproductive status”. *Apidologie* 28.3-4 (1997): 113–122.

Ito, F. “Colony composition and specialized predation on millipedes in the enigmatic ponerine ant genus *Probolomyrmex* (Hymenoptera, Formicidae)”. *Insectes Sociaux* 45.1 (1998): 79–83.

Iwata, K. “Ovarian eggs in Scarabaeoidea (Coleoptera)”. *Seibutsu Kenkyu* 10 (1966): 1–3.

Iwata, K. “The comparative anatomy of the ovary in Hymenoptera. Part I. Aculeata”. *Mushi* 29 (1955): 17–34.

Iwata, K. “The comparative anatomy of the ovary in Hymenoptera. Part II. Symphyta”. *Mushi* 31 (1958): 47–60.

Iwata, K. “The comparative anatomy of the ovary in Hymenoptera. Part IV: Proctotrupoidea and Agriotypidae (Ichneumonoidea) with descriptions of ovarian eggs”. *Entomological society of Japan* 27.1 (1959): 18–27.

Iwata, K. “The comparative anatomy of the ovary in Hymenoptera. Part V. Ichneumonidae.” *Acta Hymenopterologica* 1 (1960): 115–169.

Iwata, K. “The comparative anatomy of the ovary in Hymenoptera. Supplement of Aculeata with descriptions of ovarian eggs of certain species”. *Acta Hymenopterologica* 1 (1960): 205–211.

Iwata, K. “The comparative anatomy of the ovary in Hymenoptera. VI. Chalcidoidea with descriptions of ovarian eggs”. *Acta Hymenopterologica* 1.4 (1962): 383–391.

Iwata, K. and S. F. Sakagami. “Gigantism and dwarfism in bee eggs in relation to the modes of life, with notes on the number of ovarioles”. *Japanese Journal of Ecology* 16.1 (1966): 4–16.

Jablonska, A. and S. M. Bilinski. “Structure of ovarioles in adult queens and workers of the common wasp, *Vespula germanica* (Hymenoptera: Vespidae)”. *Folia Biologica-Krakow-* 49.3/4 (2001): 191–198.

Jackson, J. T., D. R. Tarpy, and S. E. Fahrbach. “Histological estimates of ovariole number in honey bee queens, *Apis mellifera*, reveal lack of correlation with other queen quality measures”. *Journal of Insect Science* 11 (2011): 1–11.

Jahnke, S. M., L. R. Redaelli, and L. Diefenbach. “Internal reproductive organs of *Cosmoclopius nigroannulatus* (Hemiptera: Reduviidae)”. *Brazilian Journal of Biology* 66.2A (2006): 509–512.

Jean—Louis Hemptinne, A. F., J.-L. Doucet, and J.-E. Petersen. “Optimal foraging by hoverflies (Diptera: Syrphidae) and ladybirds (Coleopteraz Coccinellidae): Mechanisms”. *European Journal of Entomology* 903 (1993): 451–455.

Jenner, W. H. “*PhD Thesis*: European parasitoids of the cherry bark tortrix: assessing the ichneumonid, *Campoplex dubitator*, as a potential classical biological control agent for North America”. Diss. Simon Fraser University, 2003.

Jervis, M. and N. Kidd. “Host-feeding strategies in hymenopteran parasitoids”. *Biological Reviews* 61.4 (1986): 395–434.

Jiang, L. and B. Hua. “Morphology and chaetotaxy of the immature stages of the scorpionfly *Panorpa liui* Hua (Mecoptera: Panorpidae) with notes on its biology”. *Journal of Natural History* 47.41-42 (2013): 2691–2705.

Kafatos, F., J. Regier, G. Mazur, M. Nadel, H. Blau, W. Petri, A. Wyman, R. Gelinas, P. Moore, M. Paul, et al. “The Eggshell of Insects: Differentiation-Specific Proteins and the Control of Their Synthesis and Accumulation During”. *Biochemical Differentiation in Insect Glands* 8 (2013): 45–145.

Kai, W. S. and I. W. Thornton. “The internal morphology of the reproductive systems of some psocid species”. *Proceedings of the Royal Entomological Society of London. Series A, General Entomology* 43.1-3 (1968): 1–12.

Karban, R. “Effects of local density on fecundity and mating speed for periodical cicadas”. *Oecologia* 51.2 (1981): 260–264.

Karlsson, M. “*Master's Thesis*: Relationship between mate guarding strategies and ovarile number in Libellulidae (Odonata)”. Diss. Halmstad University, 2007.

Karlsson, M., G. Sahlén, and K. Koch. “Continuous and stepwise oocyte production in Libellulidae (Anisoptera)”. *Odonatologica* 39.2 (2010): 107–119.

Kasap, H. and R. Crowson. “Bruchidae ve Chrysomelidae (Coleoptera) familyalarinin dicsi üreme organlari”. *Türkiye Entomoloji Derneği* 4.2 (1980): 85–102.

Kaufmann, T. “Ecological and biological studies on the West African firefly *Luciola discicollis* (Coleoptera: Lampyridae)”. *Annals of The Entomological Society of America* 58.4 (1965): 414–426.

Kaufmann, T. “Ecology, biology and gonad morphology of *Gerris rufoscutellatus* (Hemiptera: Gerridae) in Fairbanks, Alaska”. *American Midland Naturalist* 86.2 (1971): 407–416.

Kaur, A., B. K. Rao, S. Thakur, and S. Raja. “Factors inducing oocyte resorption in *Leptocoris coimbatorensis* Gross (Hemiptera: Coreidae)”. *Journal of The Kansas Entomological Society* 60.3 (1987): 353–360.

Kenis, M. and N. Mills. “Evidence for the occurrence of sibling species in *Eubazus* spp. (Hymenoptera: Braconidae), parasitoids of *Pissodes* spp. weevils (Coleoptera: Curculionidae)”. *Bulletin of Entomological Research* 88.2 (1998): 149–163.

Klemperer, H. “Life history and parental behaviour of a dung beetle from neotropical rainforest, *Copris laeviceps* (Coleoptera, Scarabaeidae)”. *Journal of Zoology* 209.3 (1986): 319–326.

Klemperer, H. “Subsocial behaviour in *Oniticellus cinctus* (Coleoptera, Scarabaeidae): effect of the brood on parental care and oviposition”. *Physiological Entomology* 8.4 (1983): 393–402.

Kobayashi, Y. “Ovarian Structure of a Zeuglopteran Moth, *Neomicropteryx nipponensis* Issiki (Lepidoptera, Micropterigidae)”. *Japanese Journal of Entomology62* 62.1 (1994): 93–100.

Koçakoğlu, N. Ö., S. Candan, and Z. Suludere. “Notes on the morphology and histology of the ovarioles of *Gerris lacustris* (L.) (water strider) (Insecta: Hemiptera: Heteroptera: Gerridae)”. *Zoologischer Anzeiger* 278 (2019): 84–89.

Koch, K., M. Quast, and G. Sahln. “Morphological differences in the ovary of Libellulidae (Odonata)”. *International Journal of Odonatology* 12.1 (2009): 147–156.

Koi, S. and J. Daniels. "New and revised life history of the Florida hairstreak *Eumaeus atala* (Lepidoptera: Lycaenidae) with notes on its current conservation status". *Florida Entomologist* 98.4 (2015): 1134–1148.

Konagaya, T., N. Mutoh, M. Suzuki, R. Rutowski, and M. Watanabe. "Estimates of female lifetime fecundity and changes in the number and types of sperm stored with age and time since mating in the monandrous swallowtail butterfly, *Battus philenor* (Lepidoptera: Papilionidae) in the Arizona desert". *Applied Entomology and Zoology* 50.3 (2015): 311–316.

Koteja, J., G. Pyka-Fosciak, M. Vogelgesang, and T. Szklarzewicz. "Structure of the ovary in *Steingelia* (Sternorrhyncha: Coccinea), and its phylogenetic implications". *Arthropod Structure and Development* 32.2-3 (2003): 247–256.

Krüger, K. and N. Mills. "Observations on the biology of three parasitoids of the spruce bark beetle, *Ips typographus* (Col., Scolytidae): *Coeloides bostrychorum*, *Dendrosoter middendorffii* (Hym., Braconidae) and *Rhopalicus tutela* (Hym., Pteromalidae)". *Journal of Applied Entomology* 110.1-5 (1990): 281–291.

Kudo, S.-i., T. Nakahira, and Y. Saito. "Morphology of trophic eggs and ovarian dynamics in the subsocial bug *Adomerus triguttulus* (Heteroptera: Cydnidae)". *Canadian Journal of Zoology* 84.5 (2006): 723–728.

Kugler, J. and Y. Nitzan. "Biology of *Clausicella suturata* [Dipt.: Tachinidae] a parasite of *Ectomyelois ceratoniae* [Lep.: Phycitidae]". *Entomophaga* 22.1 (1977): 93–105.

Kugler, J., T. Orion, and J. Ishay. "The number of ovarioles in the Vespinae (Hymenoptera)". *Insectes Sociaux* 23.4 (1976): 525–533.

Kugler, J. and Z. Wollberg. "Biology of *Agrothereutes tunetanus* Haber.[Hym. Ichneumonidae] an ectoparasite of *Orgyia dubia* Tausch.[Lep. Lymantriidae]". *Entomophaga* 12.4 (1967): 363–379.

Kulshrestha, S. K. "Histology of the ovarioles and the role of nurse cells in corpus luteum formation in *Philosamia cynthia* ricini (Boisd.) (Saturniidae: Lepidoptera)". *Journal of Natural History* 4.2 (1970): 189–197.

Kulshrestha, S. K. "Observations on the ovulation and oviposition with reference to corpus luteum formation in *Musca domestica nebulo* Fabr. (Muscidae: Diptera)". *Journal of Natural History* 3.4 (1969): 561–570.

Kumar, R. "Anatomy and relationships of Thaumastocoridae (Hemiptera: Cimicoidea)". *Australian Journal of Entomology* 3.1 (1964): 48–51.

Kumar, R. "Morphology of the reproductive and alimentary systems of the Aradoidea (Hemiptera), with comments on relationships within the superfamily". *Annals of The Entomological Society of America* 60.1 (1967): 17–25.

Kuniata, L. and G. Young. "The biology of *Lepidiota reuleauxi* Brenske (Coleoptera: Scarabaeidae), a pest of sugarcane in Papua New Guinea". *Australian Journal of Entomology* 31.4 (1992): 339–343.

Küpper, S., K.-D. Klass, G. Uhl, and M. Eberhard. "Comparative morphology of the internal female genitalia in two species of Mantophasmatodea". *Zoomorphology* 138.1 (2019): 73–83.

Kuznetsova, V. G., S. Grozeva, J.-A. N. Sewlal, and S. Nokkala. "Cytogenetic characterization of the Trinidad endemic, *Arachnocoris trinitatus* Bergroth: the first data for the tribe Arachnocorini (Heteroptera: Cimicomorpha: Nabidae)". *Folia Biologica* 55.1-2 (2007): 17–26.

Kuznetsova, V. G., S. Nokkala, and D. E. Shcherbakov. "Karyotype, reproductive organs, and pattern of gametogenesis in *Zorotypus hubbardi* Caudell (Insecta: Zoraptera, Zorotypidae), with discussion on relationships of the order". *Canadian Journal of Zoology* 80.6 (2002): 1047–1054.

LaChance, L. E. and S. B. Bruns. "Oogenesis and radiosensitivity in *Cochliomyia hominivorax* (Diptera: Calliphoridae)". *The Biological Bulletin* 124.1 (1963): 65–83.

Lachmann, A. "Sexual receptivity and post-emergence ovarian development in females of *Coproica vagans* (Diptera: Sphaeroceridae)". *Physiological Entomology* 23.4 (1998): 360–368.

Lal, K. "The biology of Scottish Psyllidae". *Transactions of The Royal Entomological Society of London* 82.2 (1934): 363–385.

Launois-Luong, M.-H. and M. Lecoq. "Sexual maturation and ovarian activity in *Rhammatocerus schistocercoides* (Orthoptera: Acrididae), a pest grasshopper in the state of Mato Grosso in Brazil". *Environmental Entomology* 25.5 (1996): 1045–1051.

Lauziere, I., J. C. Legaspi, B. C. Legaspi, J. W. Smith, and W. A. Jones. "Life-history studies of *Lydella jalisco* (Diptera: Tachinidae), a parasitoid of *Eoreuma loftini* (Lepidoptera: Pyralidae)". *BioControl* 46.1 (2001): 71–90.

Laws, A. and A. Joern. "Variable effects of dipteran parasitoids and management treatment on grasshopper fecundity in a tallgrass prairie". *Bulletin of Entomological Research* 102.2 (2012): 123–130.

Leather, S., P. W. Wellings, and K. Walters. "Variation in ovariole number within the Aphidoidea". *Journal of Natural History* 22.2 (1988): 381–393.

Lemos, W. d. P., F. d. S. Ramalho, J. E. Serrão, and J. C. Zanuncio. "Morphology of female reproductive tract of the predator *Podisus nigrispinus* (Dallas) (Heteroptera: Pentatomidae) fed on different diets". *Brazilian Archives of Biology and Technology* 48.1 (2005): 129–138.

Lemos, W., J. Zanuncio, F. Ramalho, V. Zanuncio, and J. Serrão. "Herbivory affects ovarian development in the zoophytophagous predator *Brontocoris tabidus* (Heteroptera, Pentatomidae)". *Journal of Pest Science* 83.2 (2010): 69–76.

Leprince, D. and L. Foil. "Relationships among body size, blood meal size, egg volume, and egg production of *Tabanus fuscicostatus* (Diptera: Tabanidae)". *Journal of Medical Entomology* 30.5 (1993): 865–871.

Leprince, D. and D. Lewis. "Aspects of the biology of female *Chrysops univittatus* (Diptera: Tabanidae) in southwestern Quebec". *The Canadian Entomologist* 115.4 (1983): 421–425.

Lisboa, L., J. Serrão, C. Cruz-Landim, and L. Campos. "Effect of larval food amount on ovariole development in queens of *Trigona spinipes* (Hymenoptera, Apinae)". *Anatomia, Histologia, Embryologia* 34.3 (2005): 179–184.

Liu, W.-X., W. Wang, L.-S. Cheng, J.-Y. Guo, and F.-H. Wan. "Contrasting patterns of ovarian development and oogenesis in two sympatric host-feeding parasitoids, *Diglyphus isaea* and *Neochrysocharis formosa* (Hymenoptera: Eulophidae)". *Applied Entomology and Zoology* 49.2 (2014): 305–314.

Livingstone, D. and M. Yacoob. "Female accessory glands and sperm reception in Tingidae (Heteroptera)". *Proceedings: Animal Sciences* 99.6 (1990): 431–446.

Llorente-Bousquets, J., S. Nieves-Uribe, A. Flores-Gallardo, B. C. Hernández-Mejia, and J. Castro-Gerardino. "Chorionic sculpture of eggs in the subfamily Dismorphiinae (Lepidoptera: Papilionoidea: Pieridae)". *Zootaxa* 2018.4429 (2018): 201–246.

Loan, C. and F. Holdaway. "*Microctonus aethiops* (Nees) auctt. and *Perilitus rutilus* (Nees) (Hymenoptera: Braconidae), European parasites of *Sitona* weevils (Coleoptera: Curculionidae)". *The Canadian Entomologist* 93.12 (1961): 1057–1079.

Louis, D. and R. Kumar. "Morphology of the alimentary and reproductive organs in Reduviidae (Hemiptera: Heteroptera) with comments on interrelationships within the family". *Annals of The Entomological Society of America* 66.3 (1973): 635–639.

Luft, P. A. "Experience affects oviposition in *Goniozus nigrifemur* (Hymenoptera: Bethylidae)". *Annals of The Entomological Society of America* 86.4 (1993): 497–505.

Lumbreras, C., E. Galante, and J. Mena. "Ovarian condition as an indicator of the phenology of *Bubas bubalus* (Coleoptera: Scarabaeidae)". *Annals of The Entomological Society of America* 84.2 (1991): 190–194.

Ma, N., L. Cai, and B. Hua. "Comparative morphology of the eggs in some Panorpidae (Mecoptera) and their systematic implication". *Systematics and Biodiversity* 7.4 (2009): 403–417.

Ma, N. and B. Hua. "Fine structure and formation of the eggshell in scorpionfly *Panorpa liui* Hua (Mecoptera: Panorpidae)". *Microscopy Research and Technique* 72.7 (2009): 495–500.

Ma, W. and S. Ramaswamy. "Histological changes during ovarian maturation in the tarnished plant bug, *Lygus lineolaris* (Palisot de Beauvois) (Hemiptera: Miridae)". *International Journal of Insect Morphology and Embryology* 16.5-6 (1987): 309–322.

MacDonald, J. F. and R. Matthews. "Nesting biology of the southern yellowjacket, *Vespula squamosa* (Hymenoptera: Vespidae): social parasitism and independent founding". *Journal of The Kansas Entomological Society* 57.1 (1984): 134–151.

Makert, G. R., R. J. Paxton, and K. Hartfelder. “Ovariole number—a predictor of differential reproductive success among worker subfamilies in queenless honeybee (*Apis mellifera* L.) colonies”. *Behavioral Ecology and Sociobiology* 60.6 (2006): 815–825.

Mangan, R. L. “Reproductive behavior of the cactus fly, *Odontoloxozus longicornis*, male territoriality and female guarding as adaptive strategies”. *Behavioral Ecology and Sociobiology* 4.3 (1979): 265–278.

Marchini, D., G. D. Bene, and R. Dallai. “Functional morphology of the female reproductive apparatus of *Stephanitis pyrioides* (Heteroptera, Tingidae): a novel role for the pseudospermathecae”. *Journal of Morphology* 271.4 (2010): 473–482.

Martínez, I. and M. Cruz. “Effects of nourishment on the gonadal maturation in *Canthon cyanellus* cyanellus LeConte (Coleoptera: Scarabaeidae: Scarabaeinae)”. *The Coleopterists' Bulletin* 52.3 (1998): 237–244.

Martínez, I. and C. Huerta. “Coordinated activity of the ovary, pars intercerebralis and corpus allatum during the prenesting and nesting cycles of *Copris incertus* Say (Coleoptera Scarabaeidae: Scarabaeinae)”. *The Coleopterists' Bulletin* 51.4 (1997): 351–363.

Martins, G. F. and J. E. Serrão. “A comparative study of the ovaries in some Brazilian bees (Hymenoptera; Apoidea)”. *Papeis Avulsos de Zoologia (São Paulo)* 44.3 (2004): 45–53.

Mason, L., S. Johnson, and J. Woodring. “Seasonal and ontogenetic examination of the reproductive biology of *Pseudoplusia includens* (Lepidoptera: Noctuidae)”. *Environmental Entomology* 18.6 (1989): 980–985.

Mason, P. C. “*PhD Thesis*: Alpine grasshoppers (Orthoptera: Acrididae) in the southern alps of Canterbury, New Zealand”. Diss. University of Canterbury. Zoology, 1971.

Masuko, K. “Thelytokous parthenogenesis in the ant *Strumigenys hexamera* (Hymenoptera: Formicidae)”. *Annals of The Entomological Society of America* 106.4 (2013): 479–484.

Matesco, V. C., C. F. Schwertner, and J. Grazia. “Morphology of the immatures and biology of *Chinavia longicorialis* (Breddin) (Hemiptera: Pentatomidae)”. *Neotropical Entomology* 38.1 (2009): 74–82.

Mateus, S., F. B. Noll, and R. Zucchi. “Morphological caste differences in the neotropical swarm-founding polistine wasps: *Parachartergus smithii* (Hymenoptera: Vespidae)”. *Journal of The New York Entomological Society* 62.4 (1997): 129–139.

Mathai, S., V. Nair, et al. “Histomorphological changes induced in the ovary of *Spodoptera mauritia* Boisd. (Lepidoptera: Noctuidae) by treatment with a juvenile hormone analogue”. *Proceedings of The National Academy of Sciences, India Section B: Biological Sciences* 856.3 (1990): 253–258.

Matsuzaki, M. “Electron microscopic studies on the oogenesis of dragonfly and cricket with special reference to the panoistic ovaries”. *Development, Growth and Differentiation* 13.4 (1971): 379–398.

Matsuzaki, M. and H. Ando. “Ovarian structures of the adult alderfly, *Sialis mitsuhashii* Okamoto (Megaloptera: Sialidae)”. *International Journal of Insect Morphology and Embryology* 6.1 (1977): 17–29.

Mazurkiewicz-Kania, M. “Differentiation of follicular cells in polytrophic ovaries of Pieridae and Nymphalidae (Insecta: Lepidoptera)”. *Acta Biologica Cracoviensia. Series Botanica. Supplement* 52.1 (2010): 1–98.

Mazurkiewicz-Kania, M., B. Simiczyjew, and I. Jkedrzejowska. “Differentiation of follicular epithelium in polytrophic ovaries of *Pieris napi* (Lepidoptera: Pieridae)—how far to *Drosophila* model”. *Protoplasma* 256.5 (2019): 1–15.

McLintock, J. and K. Depner. “A review of the life-history and habits of the horn fly, *Siphona irritans* (L.) (Diptera: Muscidae)”. *The Canadian Entomologist* 86.1 (1954): 20–33.

Meats, A., H. Holmes, and G. Kelly. “Laboratory adaptation of *Bactrocera tryoni* (Diptera: Tephritidae) decreases mating age and increases protein consumption and number of eggs produced per milligram of protein”. *Bulletin of Entomological Research* 94.6 (2004): 517–524.

Meier, R., M. Kotrba, and P. Ferrar. “Ovoviviparity and viviparity in the Diptera”. *Biological Reviews* 74.3 (1999): 199–258.

Melo, G. a., J. G. Rozen, et al. “Biology and immature stages of the bee tribe Tetrapediini (Hymenoptera: Apidae)”. *American Museum Novitates* 2002.3377 (2002): 1–45.

Mendes, L. “Sur deux nouvelles Nicoletiidae (Zygentoma) cavernicoles de Grèce et de Turquie et remarques sur la systématique de la famille”. *Revue Suisse de Zoologie* 95.3 (1988): 751–772.

Mercer, C. and P. King. “Ovarian development in black beetle, *Heteronychus arator* (Coleoptera: Scarabaeidae)”. *New Zealand Entomologist* 6.2 (1976): 165–170.

Messina, F. J. “Comparative biology of the goldenrod leaf beetles, *Trirhabda virgata* Leconte and *T. borealis* Blake (Coleoptera: Chrysomelidae)”. *The Coleopterists' Bulletin* 36.2 (1982): 255–269.

Michalik, A., M. Kalandyk-Kołodziejczyk, E. Simon, M. Kobiałka, and T. Szklarzewicz. “Ovaries of *Puto superbus* and *Ceroputo pilosellae* (Hemiptera: Coccoidea): Morphology, ultrastructure, phylogenetic and taxonomic implications”. *European Journal of Entomology* 110.3 (2013): 527–534.

Mizumoto, M. and F. Nakasuji. “Egg size manipulation in the migrant skipper, *Parnara guttata guttata* (Lepidoptera: Hesperiidae), in response to different host plants”. *Population Ecology* 49.2 (2007): 135–140.

Murad, H. and M. S. Ansari. “Histomorphology of the Female Reproductive Organ of the Cattle Fly, *Hippobosca maculata* Lch (Diptera: Hippoboscidae)”. *Netherlands Journal of Zoology* 31.2 (1980): 466–471.

Nalepa, C., K. Kidd, and K. Ahlstrom. “Biology of *Harmonia axyridis* (Coleoptera: Coccinellidae) in winter aggregations”. *Annals of The Entomological Society of America* 89.5 (1996): 681–685.

Nandchahal, N. “Reproductive organs of *Gryllodes sigillatus* (Walker) (Orthoptera: Gryllidae)”. *Journal of Natural History* 6.2 (1972): 125–131.

Nealis, V. and S. Fraser. “Rate of development, reproduction, and mass-rearing of *Apanteles fumiferanae* Vier. (Hymenoptera: Braconidae) under controlled conditions”. *The Canadian Entomologist* 120.3 (1988): 197–204.

Neog, K., B. G. Unni, S. Dey, C. Z. Renthlei, S. E. Reddy, P. Dutta, P. Sonowal, and R. K. Rajan. “Studies on the endocrine regulation of reproduction and ultra structure of brain and reproductive organs of muga silkworm *Antheraea assamensis*, Helfer (Lepidoptera: Saturniidae)”. *World Journal of Pharmacy and Pharmaceutical Sciences* 3.1 (2014): 1–26.

New, T. “Communal oviposition and egg-brooding in a psocid, *Peripsocus nitens* (Insecta: Psocoptera) in Chile”. *Journal of Natural History* 19.3 (1985): 419–423.

New, T. “Ovariolar dimorphism and repagula formation in some South American Ascalaphidae (Neuroptera)”. *Journal of Entomology Series A, General Entomology* 46.1 (1971): 73–77.

Ngernsiri, L., W. Piyajaraprasert, W. Wisoram, and D. J. Merritt. “Structure of the female reproductive system of the lac insect, *Kerria chinensis* (S ternorrhyncha, C occoidea: K erridae)”. *Acta Zoologica* 96.3 (2015): 312–318.

Nokkala, S. “Cytological characteristics of chromosome behaviour during female meiosis in *Sphinx ligustri* L. (Sphingidae, Lepidoptera)”. *Hereditas* 106.2 (1987): 169–179.

Noll, F. B., J. W. Wenzel, and R. Zucchi. “Evolution of caste in Neotropical swarm-founding wasps (Hymenoptera: Vespidae; Epiponini)”. *American Museum Novitates* 2004.3467 (2004): 1–24.

O'Connell, C. V. “*PhD Thesis*: A study of the internal anatomy of *Acanthocephala thomasi* Uhler (Hemiptera, Coreidae)”. Diss. The University of Arizona, 1959.

O'Neill, K. M., C. M. Delphia, and R. P. O'Neill. “Oocyte size, egg index, and body lipid content in relation to body size in the solitary bee *Megachile rotundata*”. *PeerJ* 2 (2014): 1–15.

Obata, S. “Mating refusal and its significance in females of the ladybird beetle, *Harmonia axyridis*”. *Physiological Entomology* 13.2 (1988): 193–199.

Oberhauser, K. S. “Fecundity, lifespan and egg mass in butterflies: effects of male-derived nutrients and female size”. *Functional Ecology* 11.2 (1997): 166–175.

Odendaal, F. J. “Mature egg number influences the behavior of female *Battus philenor* butterflies”. *Journal of Insect Behavior* 2.1 (1989): 15–25.

Ogorzałek, A. and A. Trochimczuk. “Ovary structure in a presocial insect, *Elasmucha grisea* (Heteroptera, Acanthosomatidae)”. *Arthropod Structure and Development* 38.6 (2009): 509–519.

Oguchi, K., H. Shimoji, Y. Hayashi, and T. Miura. “Reproductive organ development along the caste differentiation pathways in the dampwood termite *Hodotermopsis sjostedti*”. *Insectes Sociaux* 63.4 (2016): 519–529.

Ohl, M. and D. Linde. “Ovaries, ovarioles, and oocytes in apoid wasps, with special reference to cleptoparasitic species (Hymenoptera: Apoidea: Sphecidae)”. *Journal of The Kansas Entomological Society* 76.2 (2003): 147–159.

Olton, G. and E. Legner. “Biology of *Tachinaephagus zealandicus* (Hymenoptera: Encyrtidae), parasitoid of synanthropic Diptera”. *The Canadian Entomologist* 106.8 (1974): 785–800.

Osbrink, W. L. and M. K. Rust. “Fecundity and longevity of the adult cat flea, *Ctenocephalides felis* felis (Siphonaptera: Pulicidae)”. *Journal of Medical Entomology* 21.6 (1984): 727–731.

Özyurt, N., S. Candan, and Z. Suludere. “The morphology and histology of the female reproductive system of *Graphosoma lineatum* (Heteroptera: Pentatomidae) based on light and scanning electron microscope studies”. *International Journal of Scientific Research* 2.12 (2013): 42–46.

Papávcek, M. and T. Soldán. “Structure and development of the reproductive system in *Aphelocheirus aestivalis* (Hemiptera: Heteroptera: Nepomorpha: Aphelocheiridae)”. *Acta Entomologica Musei Nationalis Pragae* 48.2 (2008): 299–318.

Parkash, R., V. Sharma, and B. Kalra. “Climatic adaptations of body melanisation in *Drosophila melanogaster* from Western Himalayas”. *Fly* 2.3 (2008): 111–117.

Penney, M. M. “Diapause and reproduction in *Nebria brevicollis* (F.) (Coleoptera: Carabidae)”. *The Journal of Animal Ecology* 38.1 (1969): 219–233.

Perez-Mendoza, J., J. Throne, and J. Baker. “Ovarian physiology and age-grading in the rice weevil, *Sitophilus oryzae* (Coleoptera: Curculionidae)”. *Journal of Stored Products Research* 40.2 (2004): 179–196.

Perveen, F. “Effects of sublethal doses of Chlorfluazuron on ovarioles in the common cutworm, *Spodoptera litura* (F.) (Lepidoptera: Noctuidae)”. *Journal of Life Sciences* 5.8 (2011): 609–613.

Perveen, F. and T. Miyata. “Effects of sublethal dose of chlorfluazuron on ovarian development and oogenesis in the common cutworm *Spodoptera litura* (Lepidoptera: Noctuidae)”. *Annals of The Entomological Society of America* 93.5 (2000): 1131–1137.

Petersen, W. “Beiträge zur morphologie der Lepidopteren”. *Proceedings of the Russian Academy of Sciences* 9.6 (1900): 1–144.

Phipps, J. “The structure and maturation of the ovaries in British Acrididae (Orthoptera).” *Transactions of The Royal Entomological Society of London* 100.9 (1949): 233–247.

Pijnacker, L. and M. Ferwerda. “Additional chromosome duplication in female meiotic prophase of *Sipyloidea sipylus* Westwood (Insecta, Phasmida), and its absence in male meiosis”. *Experientia* 34.12 (1978): 1558–1560.

Pisno, R. M., K. Salazar, J. Lino-Neto, J. E. Serrão, and O. DeSouza. “Termitariophily: expanding the concept of termitophily in a physogastric rove beetle (Coleoptera: Staphylinidae)”. *Ecological Entomology* 44.3 (2019): 305–314.

Polilov, A. “Anatomy of the smallest Coleoptera, featherwing beetles of the tribe Nanosellini (Coleoptera, Ptiliidae), and limits of insect miniaturization”. *Entomological Review* 88.1 (2008): 26–33.

Pollock, J. “Viviparous adaptations in the acalyptrate genera Pachylophus (Chloropidae) and Cyrtona (Curtonotidae) (Diptera: Schizophora)”. *Annals of The Natal Museum* 37.1 (1996): 183–189.

Portman, S., J. Frank, R. McSorley, and N. Leppla. “Fecundity of *Larra bicolor* (Hymenoptera: Crabronidae) and its implications in parasitoid: host interaction with mole crickets (Orthoptera: Gryllotalpidae: Scapteriscus)”. *Florida Entomologist* 92.1 (2009): 58–64.

Price, P. W. “Energy allocation in ephemeral adult insects”. *The Ohio Journal of Science* 74.6 (1974): 380–387.

Price, P. W. “Parasitiods utilizing the same host: adaptive nature of differences in size and form”. *Ecology* 53.1 (1972): 190–195.

Price, R. and H. Brown. “Reproductive performance of the African migratory locust, *Locusta migratoria* migratorioides (Orthoptera: Acrididae), in a cereal crop environment in South Africa”. *Bulletin of Entomological Research* 80.4 (1990): 465–472.

Pritsch, M. and J. Büning. “Germ cell cluster in the panoistic ovary of Thysanoptera (Insecta)”. *Zoomorphology* 108.5 (1989): 309–313.

Punacker, L. P. and J. Godeke. "Development of ovarian follicle cells of the stick insect, *Carausius morosus* Br. (Phasmatodea), in relation to their function". *International Journal of Insect Morphology and Embryology* 13.1 (1984): 21–28.

Pyka-Fosciak, G. and T. Szklarzewicz. "Germ cell cluster formation and ovariole structure in viviparous and oviparous generations of the aphid *Stomaphis quercus*". *International Journal of Developmental Biology* 52.2-3 (2003): 259–265.

Quednau, F. and H. Guevremont. "Observations on mating and oviposition behaviour of *Priopoda nigricollis* (Hymenoptera: Ichneumonidae), a parasite of the birch leaf-miner, *Fenusa pusilla* (Hymenoptera: Tenthredinidae)". *The Canadian Entomologist* 107.11 (1975): 1199–1204.

Rabeeth, M., T. Sakthivel, and S. Janarthanan. "The internal reproductive organs of Lygaeid bug, *Spilostethus pandurus* (Heteroptera: Lygaeidae)-gross morphology and histomorphology". *Journal of Entomological Research* 40.4 (2016): 347–356.

Ramaswamy, S., G. Mbata, and N. Cohen. "Necessity of juvenile hormone for choriogenesis in the moth, *Heliothis virescens* (Noctuidae)". *Invertebrate Reproduction and Development* 17.1 (1990): 57–63.

Ramírez-Cruz, A., C. Llanderal-Cázares, and R. Racotta. "Ovariole structure of the cochineal scale insect, *Dactylopius coccus*". *Journal of Insect Science* 8.20 (2008): 2–5.

Ray, A. and P. Ramamurty. "Sources of RNA supply to the oocytes in *Crynodes peregrinus* Fuessly (Coleoptera: Chrysomelidae)". *International Journal of Insect Morphology and Embryology* 8.2 (1979): 113–122.

Reed, H. C. and R. D. Akre. "Morphological comparisons between the obligate social parasite, *Vespula austriaca* (Panzer), and its host, *Vespula acadica* (Sladen) (Hymenoptera: Vespidae)". *Psyche: A Journal of Entomology* 89.1-2 (1982): 183–195.

Regis, L. "Functional compensatory hypertrophy resulting from spontaneous or induced atrophy disconnecting one of the ovaries of *Triatoma infestans* (Heteroptera, Reduviidae, Triatominae)". *Annales de Biologie Animale Biochimie Biophysique* 17.6 (1977): 961–969.

Reichardt, T. R. and T. D. Galloway. "Seasonal occurrence and reproductive status of *Opisocrostis bruneri* (Siphonaptera: Ceratophyllidae), a flea on Franklin's ground squirrel, *Spermophilus franklinii* (Rodentia: Sciuridae) near Birds Hill Park, Manitoba". *Journal of Medical Entomology* 31.1 (1994): 105–113.

Reinhardt, K., G. Köhler, S. Maas, and P. Detzel. "Low dispersal ability and habitat specificity promote extinctions in rare but not in widespread species: the Orthoptera of Germany". *Ecography* 28.5 (2005): 593–602.

Richards, K. W. "Ovarian development, ovariole number, and relationship to body size in *Psithyrus* spp. (Hymenoptera: Apidae) in Southern Alberta". *Journal of The Kansas Entomological Society* 67.2 (1994): 156–168.

Richter, P. and C. Baker. "Ovariole number in Scarabaeoidea (Coleoptera: Lucanidae, Passalidae, Scarabaeidae)". *Proceedings of The Entomological Society of Washington* 76.4 (1974): 480–498.

Riley, C. V., A. S. Packard, C. Thomas, et al. *First anuual report of the United States entomological commission for the year 1877: Relating to the Rocky Mountain locust and the best methods of preventing its injuries and of guarding against its invasions, in pursuance of an appropriation made by congress for this purpose*. US Government Printing Office, 1878.

Robertson, H. "Sperm transfer in the ant *Carebara vidua* F. Smith (Hymenoptera: Formicidae)". *Insectes Sociaux* 42.4 (1995): 411–418.

Robertson, J. "Ovariole numbers in Coleoptera". *Canadian Journal of Zoology* 39.3 (1961): 245–263.

Root, R. B. and F. J. Messina. "Defensive adaptations and natural enemies of a case-bearing beetle, *Exema canadensis* (Coleoptera: Chrysomelidae)". *Psyche: A Journal of Entomology* 90.1-2 (1983): 67–80.

Rouibah, M., A. López-López, J. J. Presa, and S. Doumandji. "A molecular phylogenetic and phylogeographic study of two forms of *Calliptamus barbarus* (Costa 1836) (Orthoptera: Acrididae, Calliptaminae) from two regions of Algeria". *Annales de la Société entomologique de France (NS)* 52.2 (2016): 77–87.

Rozen Jr, J. G. "New taxa of brachynomadine bees (Apidae, Nomadinae)". *American Museum Novitates* 1997.3200 (1997): 1–26.

Rozen Jr, J. G., A. Roig-Alsina, and B. A. Alexander. "The cleptoparasitic bee genus *Rhopalolemma*: with reference to other Nomadinae (Apidae), and biology of its host Protodufourea (Halictidae, Rophitinae)". *American Museum novitates* 1997.3194 (1997): 1–28.

Rozen, J. G. "Eggs, ovariole numbers, and modes of parasitism of cleptoparasitic bees, with emphasis on Neotropical species (Hymenoptera: Apoidea)". *American Museum Novitates* 2003.3413 (2003): 1–36.

Rozen, J. G. "Ovarian formula, mature oocyte, and egg index of the bee *Ctenoplectra* (Hymenoptera: Apoidea: Apidae)". *Journal of The Kansas Entomological Society* 76.4 (2003): 640–642.

Rozen, J. G. and H. G. Hall. "Nesting and developmental biology of the cleptoparasitic bee *Stelis ater* (Anthidiini) and its host, *Osmia chalybea* (Osmiini) (Hymenoptera: Megachilidae)". *American Museum Novitates* 2011.3707 (2011): 1–38.

Rozen, J. G. and S. M. Kamel. "Investigations on the biologies and immature stages of the cleptoparasitic bee genera *Radoszkowskiana* and *Coelioxys* and their *Megachile* hosts (Hymenoptera: Apoidea: Megachilidae: Megachilini)". *American Museum Novitates* 2007.3573 (2007): 1–43.

Rozen, J. G. and S. M. Kamel. "Last larval instar and mature oocytes of the Old World cleptoparasitic bee *Stelis murina*, including a review of *Stelis* biology (Apoidea: Megachilidae: Megachilinae: Anthidiini)". *American Museum Novitates* 2009.3666 (2009): 1–19.

Rozen, J. G., G. A. Melo, A. J. C. Aguiar, and I. Alves-dos-Santos. "Nesting biologies and immature stages of the tapinotaspidine bee genera *Monoeca* and *Lanthanomelissa* and of their osirine cleptoparasites *Protosiris* and *Parepeolus* (Hymenoptera: Apidae: Apinae)". *American Museum Novitates* 2006.3501 (2006): 1–60.

Rozen, J. G., J. Straka, and K. Rezkova. "Oocytes, larvae, and cleptoparasitic behavior of *Biastes emarginatus* (Hymenoptera: Apidae: Nomadinae: Biastini)". *American Museum Novitates* 2009.3667 (2009): 1–15.

Rozen, J. G., S. B. Vinson, R. Coville, and G. Frankie. "Biology and morphology of the immature stages of the cleptoparasitic bee *Coelioxys chichimeca* (Hymenoptera: Apoidea: Megachilidae)". *American Museum Novitates* 2010.3679 (2010): 1–26.

Rubio G, J. D., A. E. Bustillo P, L. F. Vallejo E, J. R. Acuña Z, and P. Benavides M. "Alimentary canal and reproductive tract of *Hypothenemus hampei* (Ferrari) (Coleoptera: Curculionidae, Scolytinae)". *Neotropical Entomology* 37.2 (2008): 143–151.

Rübsam, R. and J. Büning. "Germ cell proliferation and cluster behavior in ovarioles of *Sialis flavilatera* (Megaloptera: Sialidae) during larval growth". *Arthropod Structure and Development* 46.2 (2017): 246–264.

Sadeghi, H. "The relationship between oviposition preference and larval performance in an aphidophagous hover fly, *Syrphus ribesii* L. (Diptera: Syrphidae)". *Journal Agricultural Science* 4 (2002): 1–10.

Sakagami, S. F., R. Zucchi, S. Yamane, F. Noll, and J. Camargo. "Morphological caste differences in *Agelaia vicina*, the neotropical swarm-founding polistine wasp with the largest colony size among social wasps (Hymenoptera; Vespidae)". *Sociobiology* 28.2 (1996): 207–224.

Sanderson, A. R. "Cytological investigations of parthenogenesis in gall wasps (Cynipidae, Hymenoptera)". *Genetica* 77.3 (1988): 189–216.

Sands, D. "A new genus, *Acrodipsas*, for a group of Lycaenidae (Lepidoptera) previously referred to *Pseudodipsas* C. and R. Felder, with descriptions of two new species from northern Queensland". *Australian Journal of Entomology* 18.3 (1980): 251–265.

Santeshwari. "Effect of methanolic extracts from the leaves of tulsi (Ocimum sanctum) on the ovary of *Gonocephalum brachyelytra* (Kaszab),(Coleoptera: Tenebrionidae)". *The Bioscan* 7.4 (2012): 705–709.

Santos, R. S. S. d., L. R. Redaelli, L. Diefenbach, H. P. Romanowski, and H. F. Prando. "Characterization of the imaginal reproductive diapause of *Oebalus poecilus* (Dallas) (Hemiptera: Pentatomidae)". *Brazilian Journal of Biology* 63.4 (2003): 695–703.

Sato, H. and M. Imamori. "Nesting behaviour of a subsocial African ball-roller *Kheper platynotus* (Coleoptera, Scarabaeidae)". *Ecological Entomology* 12.4 (1987): 415–425.

Satoh, T. "Comparisons between two apparently distinct forms of *Camponotus nawai* Ito (Hymenoptera: Formicidae)". *Insectes Sociaux* 36.4 (1989): 277–292.

Saunders, D. “The ovulation cycle in *Glossina morsitans* Westwood (Diptera: Muscidae) and a possible method of age determination for female tsetse flies by the examination of their ovaries”. *Transactions of The Royal Entomological Society of London* 112.9 (1960): 221–238.

Scheepens, M. and M. Wysoki. “Reproductive organs of the giant looper, *Boarmia selenaria* Schiffermüller (Leopidotera: Geometridae)”. *International Journal of Insect Morphology and Embryology* 15.1-2 (1986): 73–81.

Schilder, K., J. Heinze, and B. Hölldobler. “Colony structure and reproduction in the thelytokous parthenogenetic ant *Platythyrea punctata* (F. Smith) (Hymenoptera, Formicidae)”. *Insectes Sociaux* 46.2 (1999): 150–158.

Schultner, E., E. Blanchet, C. Pagès, G. U. Lehmann, and M. Lecoq. “Development, reproductive capacity and diet of the Mediterranean grasshopper *Arcyptera brevipennis* vicheti Harz 1975 (Orthoptera: Caelifera: Acrididae: Gomphocerinae)”. *Annales de la Société entomologique de France* 48.3-4 (2012): 299–307.

Serrão, J. E., A. P. Naves, and J. C. Zanuncio. “Modifications in the oviducts of workers and queens of *Melipona quadrifasciata* anthidioides (Hymenoptera: Apidae) with different ages”. *Protoplasma* 248.4 (2011): 767–773.

Sheldon, J. K. and E. G. MacLeod. “Studies on the biology of the Chrysopidae II. The feeding behavior of the adult of *Chrysopa carnea* (Neuroptera)”. *Psyche: A Journal of Entomology* 78.1-2 (1971): 107–121.

Sheldon, J. K. and E. G. MacLeod. “Studies on the biology of the Chrysopidae IV. A field and laboratory study of the seasonal cycle of *Chrysopa carnea* Stephens in Central Illinois (Neuroptera: Chrysopidae)”. *Transactions of The American Entomological Society (1890-)* 100.4 (1974): 437–512.

Shimada, K. and K. Maekawa. “Changes in endogenous cellulase gene expression levels and reproductive characteristics of primary and secondary reproductives with colony development of the termite *Reticulitermes speratus* (Isoptera: Rhinotermitidae)”. *Journal of Insect Physiology* 56.9 (2010): 1118–1124.

Simões, M. V. “Male and female reproductive systems of *Stolas conspersa* (Germar) (Coleoptera, Chrysomelidae, Cassidinae)”. *Revista Brasileira de Entomologia* 56.1 (2012): 19–22.

Sivinski, J., K. Vulinec, and M. Aluja. “Ovipositor length in a guild of parasitoids (Hymenoptera: Braconidae) attacking *Anastrepha* spp. fruit flies (Diptera: Tephritidae) in southern Mexico”. *Annals of The Entomological Society of America* 94.6 (2001): 886–895.

Smith, D. “Ovarioles and developing eggs in grasshoppers”. *The Canadian Entomologist* 96.9 (1964): 1255–1258.

Smith, E. and E. Salkeld. “Ovary development and oviposition rates in the plum curculio, *Conotrachelus nenuphar* (Coleoptera: Curculionidae)”. *Annals of The Entomological Society of America* 57.6 (1964): 781–787.

Soares, M. A., J. D. Batista, J. C. Zanuncio, J. Lino-Neto, and J. E. Serrão. “Ovary development, egg production and oviposition for mated and virgin females of the predator *Podisus nigrispinus* (Heteroptera: Pentatomidae)”. *Acta Scientiarum. Agronomy* 33.4 (2011): 597–602.

Soldán, T. “The structure and development of the female internal reproductive system in six European species of Ephemeroptera”. *Acta Entomologica Bohemoslovaca* 76 (1979): 353–365.

Soltani-Mazouni, N. and C. Bordereau. “Changes in the cuticle, ovaries and colleterial glands during the pseudergate and neotenic molt in *Kalotermes flavicollis* (Fabr.) (Isoptera: Kalotermitidae)”. *International Journal of Insect Morphology and Embryology* 16.3-4 (1987): 221–235.

Solulu, T., S. Simpson, and J. Kathirithamby. “The effect of strepsipteran parasitism on a tettigoniid pest of oil palm in Papua New Guinea”. *Physiological Entomology* 23.4 (1998): 388–398.

Spence, J. R. “*PhD Thesis*: Microhabitat selection and regional coexistence in water-striders (Heteropetra: Gerridae)”. Diss. University of British Columbia, 1979.

Spradbery, J. “Seasonal changes in the population structure of wasp colonies (Hymenoptera: Vespidae)”. *The Journal of Animal Ecology* 40.2 (1971): 501–523.

Spradbery, J. “The biology of *Stenogaster concinna* Van der Vecht with comments on the phylogeny of Stenogastrinae (Hymenoptera: Vespidae)”. *Australian Journal of Entomology* 14.3 (1975): 309–318.

Spradbery, J. and D. Sands. “Reproductive system and terminalia of the Old World screw-worm fly, *Chrysomya bezziana* Villeneuve (Diptera: Calliphoridae)”. *International Journal of Insect Morphology and Embryology* 5.6 (1976): 409–421.

Srinivasa Rao Vattikonda, M. M. and S. Raja. “Effect of Andrographolide on ovarian development of *Papilio demoleus* L. (Lepidoptera: Papilionidae) larvae”. *International Journal of Entomology Research* 3.2 (2018): 23–27.

Stadler, B. and A. Dixon. “Ant attendance in aphids: why different degrees of myrmecophily?” *Ecological Entomology* 24.3 (1999): 363–369.

Starmer, W. T., M. Polak, S. Pitnick, S. F. McEvey, J. S. F. Barker, and L. L. Wolf. “Phylogenetic, geographical, and temporal analysis of female reproductive trade-offs in Drosophilidae”. *Evolutionary Biology* 33 (2003): 139–171.

Stay, B. “Protein uptake in the oocytes of the cecropia moth”. *The Journal of Cell Biology* 26.1 (1965): 49–62.

Stewart, L., J.-L. Hemptinne, and A. Dixon. “Reproductive tactics of ladybird beetles: relationships between egg size, ovariole number and developmental time”. *Functional Ecology* 5.3 (1991): 380–385.

Stille, B. and L. Dävring. “Meiosis and reproductive strategy in the parthenogenetic gall wasp *Diplolepis rosae* (L.) (Hymenoptera, Cynipidae)”. *Hereditas* 92.2 (1980): 353–362.

Stringer, I. “The female reproductive system of *Costelytra zealandica* (White) (Coleoptera: Scarabaeidae: Melolonthinae)”. *New Zealand Journal of Zoology* 15.4 (1988): 513–533.

Sturm, R. “Relationship between body size and reproductive capacity in females of the black field cricket (Orthoptera, Gryllidae)”. *Linzer biologische Beiträge* 48 (2016): 1–12.

Su, X. H., J. L. Chen, X. J. Zhang, W. Xue, H. Liu, and L. X. Xing. “Testicular development and modes of apoptosis during spermatogenesis in various castes of the termite *Reticulitermes labralis* (Isoptera: Rhinotermitidae)”. *Arthropod Structure and Development* 44.6 (2015): 630–638.

Susa, K. and M. Watanabe. “Egg production in *Sympetrum infuscatum* (Selys) females living in a forest-paddy field complex (Anisoptera: Libellulidae)”. *Odonatologica* 36.2 (2007): 159–170.

Sutherland, B. “Physiological age determination in female *Stomoxys calcitrans* Linnaeus (Diptera: Muscidae).” *The Onderstepoort Journal of Veterinary Research* 47.2 (1980): 83–88.

Svensson, B. G. and E. Petersson. “Sex-role reversed courtship behaviour, sexual dimorphism and nuptial gifts in the dance fly, *Empis borealis* (L.)” *Annales Zoologici Fennici* 24.4 (1987): 323–334.

Syme, P. D. “Observations on the longevity and fecundity of *Orgilus obscurator* (Hymenoptera: Braconidae) and the effects of certain foods on longevity”. *The Canadian Entomologist* 109.7 (1977): 995–1000.

Szklarzewicz, T. “Structure and development of the telotrophic ovariole in ensign scale insects (Hemiptera, Coccomorpha: Ortheziidae)”. *Tissue and Cell* 29.1 (1997): 31–38.

Szklarzewicz, T. “Oogenesis of *Nicoletia phytophila* (Zygentoma, Nicoletiidae). Preliminary studies”. *Recent Advances in Insect Embryology in Japan and Poland* (1987): 69–76.

Szklarzewicz, T., A. Jabłońska, and S. M. Biliński. “Ovaries of *Petrobius brevistylis* (Archaeognatha, Machilidae) and *Tricholepidion gertschi* (Zygentoma, Lepidotrichidae): morphology, ultrastructure and phylogenetic implications”. *Pedobiologia* 48.5-6 (2004): 477–485.

Szklarzewicz, T., M. Kalandyk-Kolodziejczyk, M. Kot, and A. Michalik. “Ovary structure and transovarial transmission of endosymbiotic microorganisms in *Marchalina hellenica* (Insecta, Hemiptera, Coccomorpha: Marchalinidae)”. *Acta Zoologica* 94.2 (2013): 184–192.

Szklarzewicz, T., A. Michalik, A. Czaja, et al. “Germ cell cluster formation and ovariole structure in *Puto albicans* and Crypticerya morrilli (Hemiptera: Coccinea). Phylogenetic implications.” *European Journal of Entomology* 107.4 (2010): 589–595.

Szklarzewicz, T., A. Michalik, M. Kalandyk-Kołodziejczyk, M. Kobiałka, and E. Simon. “Ovary of *Matsucoccus pini* (Insecta, Hemiptera, Coccinea: Matsucoccidae): morphology, ultrastructure, and phylogenetic implications”. *Microscopy Research and Technique* 77.5 (2014): 327–334.

Szklarzewicz, T., A. Wne k, and S. M. Biliński. “Structure of ovarioles in *Adelges laricis*, a representative of the primitive aphid family Adelgidae”. *Acta Zoologica* 81.4 (2000): 307–313.

Tachi, T. and H. Shima. “Molecular phylogeny of the subfamily Exoristinae (Diptera, Tachinidae), with discussions on the evolutionary history of female oviposition strategy”. *Systematic Entomology* 35.1 (2010): 148–163.

Taddei, C., M. Chicca, M. G. Maurizii, and V. Scali. “The germarium of panoistic ovarioles of *Bacillus rossius* (Insecta Phasmatodea): Larval differentiation”. *Invertebrate Reproduction and Development* 21.1 (1992): 47–56.

Talhouk, A. "Contributions to the knowledge of almond pests in East Mediterranean countries: I. Notes on *Eriogaster amygdali* Wilts. (Lepid., Lasiocampidae) with a description of a new subspecies by EP Wiltshire 1". *Zeitschrift Für Angewandte Entomologie* 78.1-4 (1975): 306–312.

Tanaka, M. "Developmental stages of egg follicles in *Parnassius glacialis* Butler (Lepidoptera, Papilionidae)". *Lepidoptera Science* 40.3 (1989): 167–181.

Tanton, M. and J. Epila. "Effects of DDT and Fenitrothion on field-collected larvae of a eucalypt-defoliating beetle, *Paropsis atomaria* Ol. I. Mortality, relative toxicity, development of treated larvae, and effect of the primary parasitoids." *Australian Journal of Zoology* 32.3 (1984): 325–336.

Tay, J.-W. and C.-Y. Lee. "Influences of pyriproxyfen on fecundity and reproduction of the pharaoh ant (Hymenoptera: Formicidae)". *Journal of Economic Entomology* 107.3 (2014): 1216–1223.

Taylor, B. J. and D. W. Whitman. "A test of three hypotheses for ovariole number determination in the grasshopper *Romalea microptera*". *Physiological Entomology* 35.3 (2010): 214–221.

Terkanian, B. "Effect of host deprivation on egg quality, egg load, and oviposition in a solitary parasitoid, *Chetogena edwardsii* (Diptera: Tachinidae)". *Journal of Insect Behavior* 6.6 (1993): 699–713.

Togashi, K. "Lifetime fecundity and female body size in *Paraglenea fortunei* (Coleoptera: Cerambycidae)". *Applied Entomology and Zoology* 42.4 (2007): 549–556.

Togashi, K., J. E. Appleby, H. Oloumi-Sadeghi, and R. B. Malek. "Age-specific survival rate and fecundity of adult *Monochamus carolinensis* (Coleoptera: Cerambycidae) under field conditions". *Applied Entomology and Zoology* 44.2 (2009): 249–256.

Togashi, K. and M. Itabashi. "Maternal size dependency of ovariole number in *Dastarcus helophoroides* (Coleoptera: Colydiidae)". *Journal of Forest Research* 10.5 (2005): 373–376.

Togashi, K. and H. Yamashita. "Effects of female body size on lifetime fecundity of *Monochamus urussovii* (Coleoptera: Cerambycidae)". *Applied Entomology and Zoology* 52.1 (2017): 79–87.

Tourneur, J.-C. "Factors affecting the egg-laying pattern of *Forficula auricularia* (dermaptera: Forficulidae) in three climatologically different zones of North America". *The Canadian Entomologist* 150.4 (2018): 511–519.

Tourneur, J.-C. "Oogenesis in the adult of the European earwig *Forficula auricularia* (dermaptera: Forficulidae)". *The Canadian Entomologist* 131.3 (1999): 323–334.

Trauner, J. and J. Büning. "Germ-cell cluster formation in the telotrophic meroistic ovary of *Tribolium castaneum* (Coleoptera, Polyphaga, Tenebrionidae) and its implication on insect phylogeny". *Development Genes and Evolution* 217.1 (2007): 13–27.

Tripp, H. A. "The biology of a hyperparasite, *Euceros frigidus* Cress. (Ichneumonidae) and description of the planidial stage". *The Canadian Entomologist* 93.1 (1961): 40–58.

TRL, B. "Karyotypes of some Lepidoptera chromosomes and changes in their holokinetic organisation as revealed by new cytological techniques". *Cytologia* 40.3-4 (1975): 713–726.

Tschinkel, W. R. "Relationship between ovariole number and spermathecal sperm count in ant queens: a new allometry". *Annals of The Entomological Society of America* 80.2 (1987): 208–211.

Tsutsumi, T., M. Matsuzaki, and K. Haga. "Formation of germ cell cluster in tubuliferan thrips (Thysanoptera)". *International Journal of Insect Morphology and Embryology* 24.3 (1995): 287–296.

Tworzydlo, W., S. M. Bilinski, P. Kovcárek, and F. Haas. "Ovaries and germline cysts and their evolution in Dermaptera (Insecta)". *Arthropod Structure and Development* 39.5 (2010): 360–368.

Tworzydlo, W., E. Kisiel, W. Jankowska, and S. M. Bilinski. "Morphology and ultrastructure of the germarium in panoistic ovarioles of a basal "apterygotous" insect, *Thermobia domestica*". *Zoology* 117.3 (2014): 200–206.

Uckan, F., E. Ergin, S. Sinan, and O. Sak. "Morphology of the reproductive tract and ovariole histology of *Apanteles galleriae* (Hymenoptera: Braconidae) reared on two host species". *Pakistan Journal of Biological Sciences* 6.16 (2003): 1389–1395.

Ueno, T. "Adult size and reproduction in the ectoparasitoid *Agrothereutes lanceolatus* Walker (Hym., Ichneumonidae)". *Journal of Applied Entomology* 123.6 (1999): 357–361.

Ueno, T. “Reproduction and host-feeding in the solitary parasitoid wasp *Pimpla nipponica* (Hymenoptera: Ichneumonidae)”. *Invertebrate Reproduction and Development* 35.3 (1999): 231–237.

Ueno, T. and T. Tanaka. “Comparative biology of six polyphagous solitary pupal endoparasitoids (Hymenoptera: Ichneumonidae): differential host suitability and sex allocation”. *Annals of The Entomological Society of America* 87.5 (1994): 592–598.

Ullman, D. E., D. M. Westcot, W. B. Hunter, and R. F. Mau. “Internal anatomy and morphology of *Frankliniella occidentalis* (Pergande) (Thysanoptera: Thripidae) with special reference to interactions between thrips and tomato spotted wilt virus”. *International Journal of Insect Morphology and Embryology* 18.5-6 (1989): 289–310.

Ullmann, S. L. “Oogenesis in *Tenebrio molitor*: histological and autoradiographical observations on pupal and adult ovaries”. *Development* 30.1 (1973): 179–217.

Uzsák, A. and C. Schal. “Differential physiological responses of the German cockroach to social interactions during the ovarian cycle”. *Journal of Experimental Biology* 215.17 (2012): 3037–3044.

Van Dijk, T. S. “The significance of the diversity in age composition of *Calathus melanocephalus* L. (Col., Carabidae) in space and time at Schiermonnikoog”. *Oecologia* 10.2 (1972): 111–136.

Van Rensburg, N. “A technique for rearing the black pine aphid, *Cinara cronartii* T and P, and some features of its biology (Homoptera: Aphididae)”. *Journal of The Entomological Society of Southern Africa* 44.2 (1981): 367–379.

Varadarasan, S. and T. Ananthakrishnan. “Biological studies on some gall thrips”. *Proceedings of The National Academy of Sciences, India Section B: Biological Sciences* 48 (1982): 35–43.

Vargas, R. I., L. Leblanc, R. Putoa, and J. C. Piñero. “Population dynamics of three *Bactrocera* spp. fruit flies (Diptera: Tephritidae) and two introduced natural enemies, *Fopius arisanus* (Sonan) and *Diachasmimorpha longicaudata* (Ashmead) (Hymenoptera: Braconidae), after an invasion by *Bactrocera dorsalis* (Hendel) in Tahiti”. *Biological Control* 60.2 (2012): 199–206.

Vianen, A. v. and J. v. Lenteren. “The parasite-host relationship between *Encarsia formosa* Gahan (Hym., Aphelinidae) and Trialeurodes vaporariorum (Westwood) (Horn., Aleyrodidae) XIV. Genetic and environmental factors influencing body-size and number of ovarioles of *Encarsia formosa*”. *Journal of Applied Entomology* 101.1-5 (1986): 321–331.

Villet, M. “Qualitative relations of egg size, egg production and colony size in some ponerine ants (Hymenoptera: Formicidae)”. *Journal of Natural History* 24.5 (1990): 1321–1331.

Wada, T., M. Kobayashi, and M. Shimazu. “Seasonal changes of the proportions of mated females in the field population of the rice leaf roller, *Cnaphalocrocis medinalis* Guené (Lepidoptera: Pyralidae)”. *Applied Entomology and Zoology* 15.1 (1980): 81–89.

Wagenhoff, E., R. Blum, and H. Delb. “Spring phenology of cockchafers, *Melolontha* spp. (Coleoptera: Scarabaeidae), in forests of south-western Germany: results of a 3-year survey on adult emergence, swarming flights, and oogenesis from 2009 to 2011”. *Journal of Forest Science* 60.4 (2014): 154–165.

Waloff, N. “Number and Development of Ovarioles of Some Acridoidea (Orthoptera) in Relation To Climate”. *Physiologia comparata et oecologia* 3.2 (1954): 370–390.

Ware, R. L., B. Yguel, and M. E. Majerus. “Effects of larval diet on female reproductive output of the European coccinellid *Adalia bipunctata* and the invasive species *Harmonia axyridis* (Coleoptera: Coccinellidae)”. *European Journal of Entomology* 105.3 (2008): 437–443.

Watanabe, M. and S. Matsu’ura. “Fecundity and oviposition in *Mortonagrion hirosei* Asahina, *M. selenion* (Ris), *Ischnura asiatica* (Brauer) and *I. senegalensis* (Rambur), coexisting in estuarine landscapes of the warm temperate zone of Japan (Zygoptera: Coenagrionidae)”. *Odonatologica* 35.2 (2006): 159–166.

Watanabe, M. “Multiple matings increase the fecundity of the yellow swallowtail butterfly, *Papilio xuthus* L., in summer generations”. *Journal of Insect Behavior* 1.1 (1988): 17–29.

Wensler, R. J. and J. Rempel. “The morphology of the male and female reproductive systems of the midge, *Chironomus plumosus* L.” *Canadian Journal of Zoology* 40.2 (1962): 199–229.

West, R. and M. Kenis. “Screening four exotic parasitoids as potential controls for the eastern hemlock looper, *Lambdina fiscellaria* fiscellaria (Guené) (Lepidoptera: Geometridae)”. *The Canadian Entomologist* 129.5 (1997): 831–841.

Weyda, F. “Female reproductive system of the first-instar nymphs of *Machilis helleri* (Verh.) (Thysanura: Machilidae) with special reference to the segmental arrangement of ovarioles in arthropods”. *International Journal of Insect Morphology and Embryology* 18.2-3 (1989): 85–96.

White, M. and N. Contreras. “Cytogenetics of the parthenogenetic grasshopper *Warramaba* (formerly *Moraba*) *virgo* and its bisexual relatives. V. Interaction of *W. virgo* and a bisexual species in geographic contact”. *Evolution* 67.4 (1979): 85–94.

Wightman, J. “Ovariole microstructure and vitellogenesis in *Lygocoris pabulinus* (L.) and other mirids (Hemiptera: Miridae)”. *Journal of Entomology Series A, General Entomology* 48.1 (1973): 103–115.

Wikars, L.-O. “Effects of forest fire and the ecology of fire-adapted insects”. Diss. Universitatis Upsaliensis Uppsala, 1997.

Wiktelius, S. and P. Chiverton. “Ovariole number and fecundity for the two emigrating generations of the bird cherry-oat aphid (Rhopalosiphum padi) in Sweden”. *Ecological Entomology* 10.3 (1985): 349–355.

Wildman, M. and R. Crewe. “Gamergate number and control over reproduction in *Pachycondyla krugeri* (Hymenoptera: Formicidae)”. *Insectes Sociaux* 35.3 (1988): 217–225.

Wilkes, A. “Sperm transfer and utilization by the arrhenotokous wasp *Dahlbominus fuscipennis* (Zett.) (Hymenoptera: Eulophidae)”. *The Canadian Entomologist* 97.6 (1965): 647–657.

Winnick, C. G., G. I. Holwell, and M. E. Herberstein. “Internal reproductive anatomy of the praying mantid *Ciulfina klassi* (Mantodea: Liturgusidae).” *Arthropod Structure and Development* 38.1 (2009): 60–69.

Wishart, G. and E. Monteith. “Trybliographa rapae (Westw.) (Hymenoptera: Cynipidae), a parasite of *Hylemya* spp. (Diptera: Anthomyiidae)”. *The Canadian Entomologist* 86.4 (1954): 145–154.

Wojcik, D. P. and D. Habeck. “Fire ant Myrmecophiles: Breeding period and ovariole number in *Myrmecaphodius excavaticollis* (Blanchard) and *Euparia castanea* Serville (Coleoptera: Scarabaeidae)”. *The Coleopterists' Bulletin* 31.4 (1977): 335–338.

Woolley, T. A. “Studies on the internal anatomy of the Box elder bug, *Leptocoris trivittatus* (Say) (Hemiptera, Coreidae)”. *Annals of The Entomological Society of America* 42.2 (1949): 203–226.

Xider, K. M. and H. M. Amin. “Ovarian Development of House Fly (*Musca domestica* L.) (Diptera: Muscidae)”. *Kurdistan Journal of Applied Research* 3.1 (2018): 45–51.

Yamauchi, H. and N. Yoshitake. “Origin and differentiation of the oocyte-nurse cell complex in the germarium of the earwig, *Anisolabis maritima* Borelli (dermaptera: Labiduridae)”. *International Journal of Insect Morphology and Embryology* 11.5-6 (1982): 293–305.

Yasumatsu, K. and A. Taketani. *Some remarks on the commonly known species of the genus* Diplolepis *Geoffroy in Japan*. 1967.

Yel, M., E. Eren, et al. “The anatomic and histologic structure of the female reproduction systems *Pieris rapae* (L.) (Lepidoptera: Pieridae).” *Türk Hijyen ve Deneysel Biyoloji Dergisi* 57.1 (2000): 25–34.

Yuan, W., W. Li, Y. Li, and K. Wu. “Combination of plant and insect eggs as food sources facilitates ovarian development in an omnivorous bug *Apolygus lucorum* (Hemiptera: Miridae)”. *Journal of Economic Entomology* 106.3 (2013): 1200–1208.

Zaviezo, T. and N. Mills. “Aspects of the biology of *Hyssopus pallidus* (Hymenoptera: Eulophidae), a parasitoid of the codling moth (Lepidoptera: Olethreutidae)”. *Environmental Entomology* 28.4 (1999): 748–754.

Zawadzka, M., W. Jankowska, and S. Biliński. “Egg shells of mallophagans and anoplurans (Insecta: Phthiraptera): morphogenesis of specialized regions and the relation to F-actin cytoskeleton of follicular cells”. *Tissue and Cell* 29.6 (1997): 665–673.

Zelazowska, M. and S. Biliński. “Distribution and transmission of endosymbiotic microorganisms in the oocytes of the pig louse, *Haematopinus suis* (L.) (Insecta: Phthiraptera)”. *Protoplasma* 209.3-4 (1999): 207–213.

Zhang, L., Z. Wu, J. Fan, G. Wang, et al. "Reproductive characteristics of female *Tetrastichus hagenowii* (Ratzeburg) (Hymenoptera: Eulophidae)." *Acta Entomologica Sinica* 53.1 (2010): 76–81.

Ziegler, R. and R. Van Antwerpen. "Lipid uptake by insect oocytes". *Insect Biochemistry and Molecular Biology* 36.4 (2006): 264–272.

Zrzavy, J. "Four chapters about the monophyly of insect 'orders': A review of recent phylogenetic contributions". *Acta Entomol Musei Nat Pragae* 48 (2008): 217–232.
