## Supplementary Methods for "Repeated loss of variation in insect ovary morphology highlights the role of developmental constraint in life-history evolution"

**References** **20**

[1] Department of Organismic and Evolutionary Biology, Harvard University, Cambridge, MA 02138, USA

[2] Smithsonian Tropical Research Institute, Panama City, Panama

[3] Department of Molecular Genetics and Cell Biology, University of Chicago, Chicago, IL 60637, USA

[4] Department of Biology, Universidad de Puerto Rico en Cayey, Cayey 00736, PR

[5] Department of Molecular & Cellular Biology, Harvard University, Cambridge, MA 02138, USA

* Corresponding author

### Contents

### 1 Gathering ovariole number records

We searched the published literature for references to insect ovariole number using a predetermined set of 131 search terms, entered into Google Scholar (scholar.google.com) between June and October of 2019. Each search term consisted of an insect taxonomic group and the words "ovariole number". This list was created to include all insect orders, many large insect families, and groups well-represented in the insect egg dataset. The list of search terms is available in the supplementary file 'ovariole_number_search_terms.tsv'.

For each search term, we evaluated all publications in the first page of results (ten publications). For 61 search terms that had a large number of informative hits, significant representation in the egg dataset, or that corresponded to very speciose groups, we evaluated an additional 20 publications. If a publication reported ovariole number for one or more insect species, we recorded the following information: (1) genus, (2) species name, when available, (3) taxonomic order, (4) sample size, when available, (5) ovariole number, and (6) additional notes (e.g. for eusocial insects, whether the observation was made in a reproductive or non-reproductive individual). This dataset is made publically available at Dryad (doi:10.5061/dryad.59zw3r253).

Ovariole number was recorded as either an average with deviations, a range, or a single total value. When multiple types of data were available from a single publication, we recorded only a single type, with priority given to averages over ranges, and to both over single total values. Ovariole number was recorded as the total number of ovarioles per female, summing over both the left and right adult ovaries. When authors reported ovariole number from a single ovary, the total value was calculated by doubling the reported value. When authors described differences between the two ovaries, this information was recorded in an additional notes column.

Using this approach, we gathered 3355 records for ovariole number from 460 publications. A full list of publications is provided in the supplementary file 'ovariole_number_bibliography.pdf'. We matched the scientific names to additional taxonomic information using the software TaxReformer[1] and found additional taxonomic data for 3252 of the 3355 records. We verified that TaxReformer had found a valid match by comparing the originally recorded taxonomic order to the order populated by online databases, and removed 22 taxonomic records for which these values did not match. For all subsequent analyses, we also excluded observations made in non-reproductive individuals from eusocial species (workers), as well as two observations which represented significant outliers and could not be validated using additional sources or figures[2,3].

### 2 Phylogenetic trees

The analyses herein were performed using the insect phylogeny published in Church et al, 2019[4], unless otherwise specified. This phylogeny was constructed by combining ribosomal genetic data from 1726 insect genera, published originally in the SILVA database[5], with constrained, time-calibrated nodes for each insect order, published originally in Misof et al, 2014[6]. This phylogeny is enriched for insect genera with records in the egg trait dataset, and also has considerable overlap with the genera included in this ovariole number dataset (508 genera). For generalized least squares analyses and trait model comparisons, analyses were performed over a posterior distribution of trees associated with this published phylogeny[4].

Analyses of insect family-level ovariole number, egg size, and body size were performed using the insect phylogeny published in Rainford et. al, 2014[7].

Analyses of Drosophilidae ovariole number, egg size, and body size were performed using a phylogeny newly assembled for this study. Published genetic data for 317 Drosophilidae species were retrieved from NCBI in June of 2019[8–16]. These data encompassed 41 gene regions including mitochondrial, nuclear, and ribosomal genes. When multiple sequences for a gene region were available from the same species, the one with the least amount of missing data was selected. Each gene region was aligned using the program MAFFT[17], model auto selected). Alignments were concatenated and trimmed to 3% occupancy across species using the program phyutlity[18]. Documentation including accession numbers, sequence files, and alignments are available in the supplementary directory 'https://github.com/shchurch/insect_ovariole_number_evolution_2020/phylogeny/Drosophilidae_sequences/'.

To the extent possible, sequence data were not curated beyond what was downloaded from NCBI, with the following exceptions: [1] two sequences labeled as 16S that did not align to other 16S sequences were removed manually. [2] COI sequences were trimmed to remove regions with large quantities of missing sites prior to alignment. [3] One species name (*D. albovittata*) was corrected for typographical error. [4] Sequences identified as *Drosophila crassifemur* were taxonomically corrected to *Scaptomyza crassifemur*[19].

Phylogenetic estimation of the Drosophilidae data were performed using RAxML (model GTRGAMMA), setting the split between Hawaiian *Drosophila* and *Scaptomyza* as the root of the tree[8,11]. The final tree was pruned to remove undescribed species (e.g. *Drosophila* nr *dorsigera*), and was time-calibrated using the R package ape, function chronos (default parameters, version 5.4.1)[20]. This tree is available in the supplementary file 'https://github.com/shchurch/insect_ovariole_number_evolution_2020/phylogeny/Drosophilidae_time_calibrated.tre'.

### 3 Phylogenetic regressions

#### 3.1 Combining datasets

We combined the data we collected on total ovariole number with existing datasets of egg size and shape[21], insect lifetime fecundity and dry adult body mass[22], average adult body length per insect family[23], and several lineage-specific measures of adult body size[24–28].

Ovariole number and egg size[4] data were combined by matching records at the species level (Fig 2a). When multiple records existed for a given species, the dataset was randomly shuffled and a single matching record was selected. This variation across records for the same species was accounted for in regressions by reshuffling and matching records at each iteration of the analysis. We also matched records at the genus level following the same reshuffling method, which allowed us to test whether results were robust with a larger sample size when an exact species match was not available (Fig. S1).

Average adult body length per insect family[23] was matched to the average ovariole number for the corresponding families (Fig. S3). The Rainford et al, 2016[23] dataset contains a small number of average adult body lengths at the order level (e.g. Strepsiptera), which were matched to their equivalent group in the ovariole number dataset. To test the effect of uncertainty in the estimated average ovariole number on our results, the dataset for each family was downsampled by half at each iteration of the regression analysis.

Ovariole number, egg volume, lifetime fecundity, and adult body mass[22] were combined by matching records at the species level and genus level, using the same method as described above (Figs. 2b, 3, and S2). We excluded one value from this dataset which appeared to include a typographical error for lifetime fecundity (Hymenoptera: Trichogrammatidae, lifetime fecundity recorded as 0.1[22]).

Several lineage-specific measurements for body size were matched to the ovariole number and egg size datasets, as follows: Drosophilidae thorax length[27] was matched at the species level (Fig. 2c), Orthoptera body length[26] was matched at the genus level (Fig. 2d), Hymenoptera mesosoma width[25] was matched at the genus level, and Curculonoidea elytra length[24] was matched at the genus level (Fig. S5).

#### 3.2 Phylogenetic Generalized Least Squares (PGLS) analyses

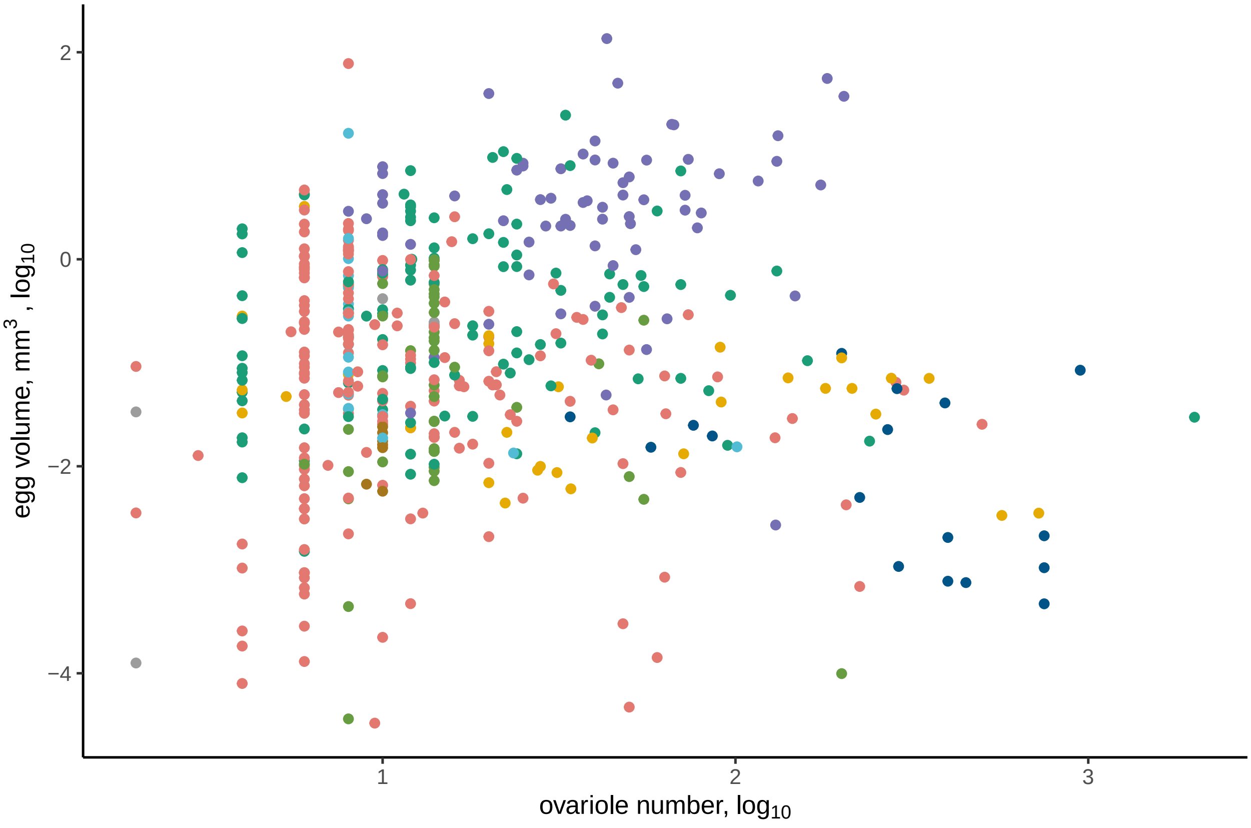

Figure S1: **Egg volume vs ovariole number, matching records at the genus level.**

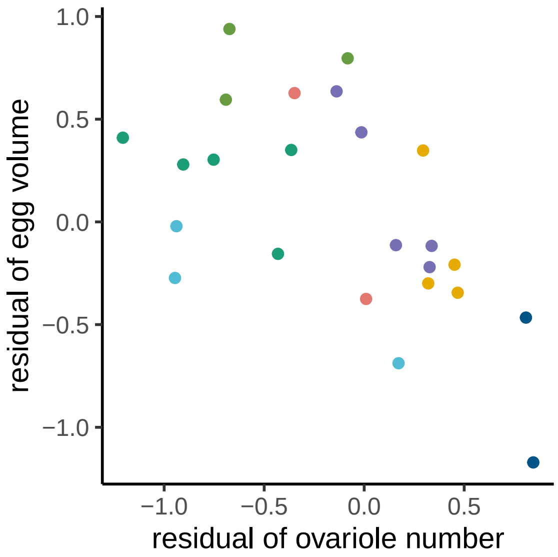

Figure S2: **Egg volume vs ovariole number, phylogenetic residuals to dry adult body mass, matching records at the species level.**

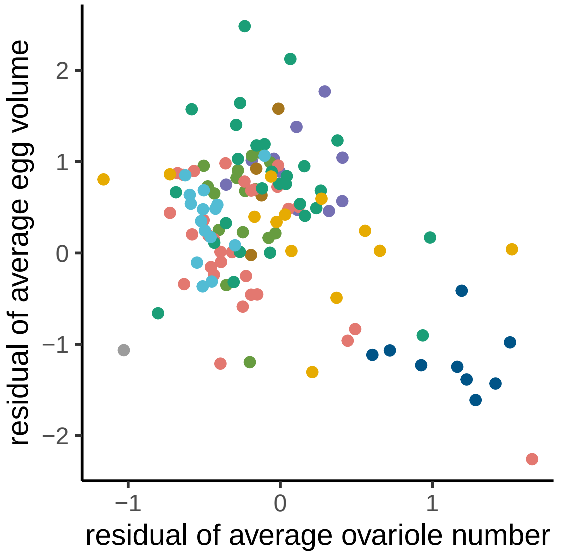

Figure S3: **Family average egg volume vs ovariole number, phylogenetic residuals to adult body length.**

We ran each Phylogenetic Generalized Least Squares (PGLS) regression over the Maximum Clade Credibilty (MCC) tree. For each regression, we also repeated each PGLS analysis 1000 times, accounting for phylogenetic and phenotypic uncertainty, using the R packages ape (version 5.4.1)[20] and nlme (version 3.1.151)[29]. In these analyses we used a Brownian Motion based covariance matrix for traits.

For regressions at the species and genus level, we reshuffled and matched records at each iteration to account for variation across records for the same taxon. For regressions at the family level we recalculated the average ovariole number per insect family, downsampling the representation for each family by half. No posterior distribution was available with the previously published family level phylogeny[7].

To account for body size, we calculated the phylogenetic residuals[30] of each trait to body size, and then compared the evolution of these residuals using a PGLS regression.

For regressions of egg size and ovariole number when accounting for adult body size, we compared the results of our regression analyses to distributions estimated using simulated data under alternative hypotheses. We fit a Brownian motion model to the phylogenetic residuals of egg size and body size (R package geiger, version 2.0.7)[31], and then used the parameters of this fitted model to simulate new datasets (R package phylolm, version 2.6.2)[32]. We performed this resimulation using the datasets of egg size and body length at the family level, and egg size and body mass at the genus level. We simulated 1000 datasets each under two hypotheses: no correlation (slope=0) and a strong negative correlation (slope=-1). We performed the regressions as described above and compared the distribution of p-values and slopes to values from regressions on observed data.

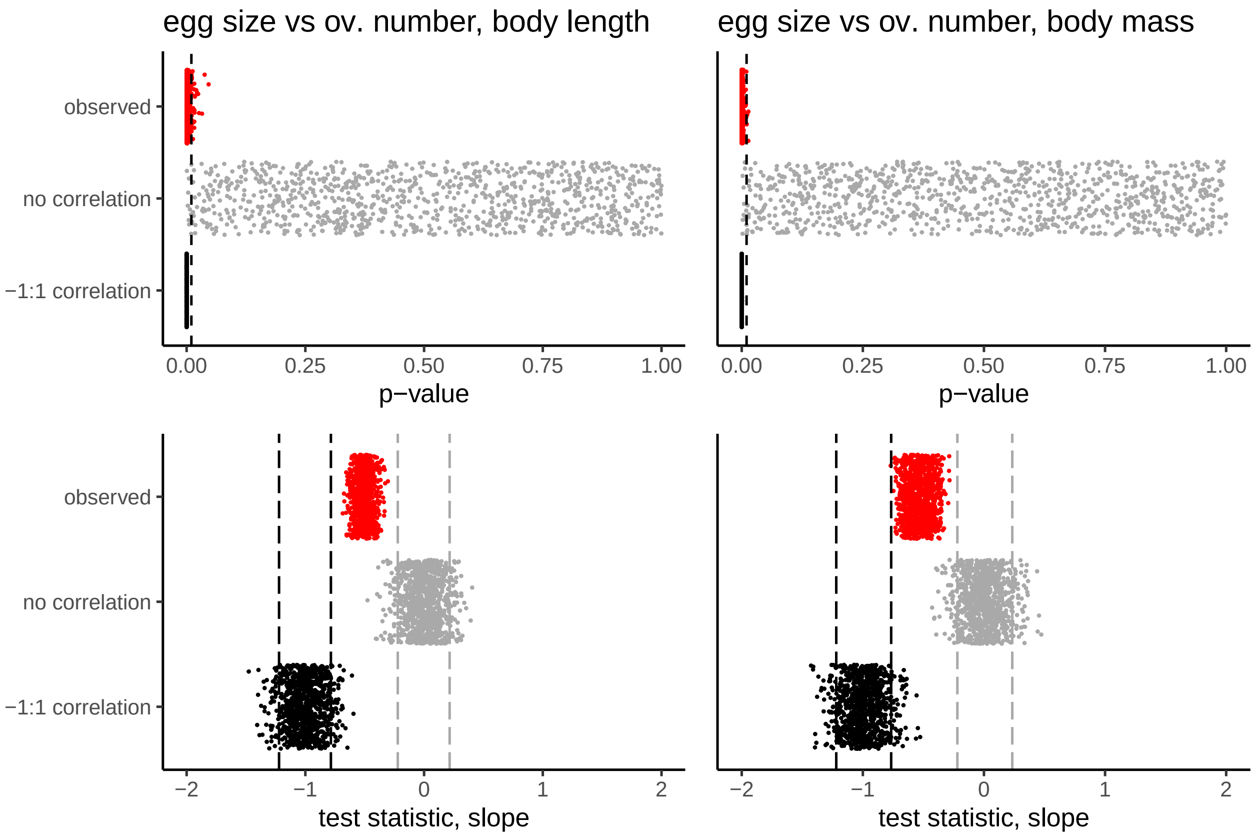

Figure S4: **Using simulated data to test alternative hypotheses of evolutionary relationships.** Top row, distributions of p-values over 1000 replicate regressions, dashed black line indicates threshold of 0.01. Bottom row, distribution of estimated slopes between egg size and ovariole number, dashed lines indicate 95% interval of simulated distributions. Left, comparing egg size and ovariole number, accounting for body length at the family level. Right, comparing egg size and ovariole number, accounting for body mass at the genus level. Red=observed values, gray=simulated with no correlation, black=simulated with a -1:1 correlation. n=1000 regressions.

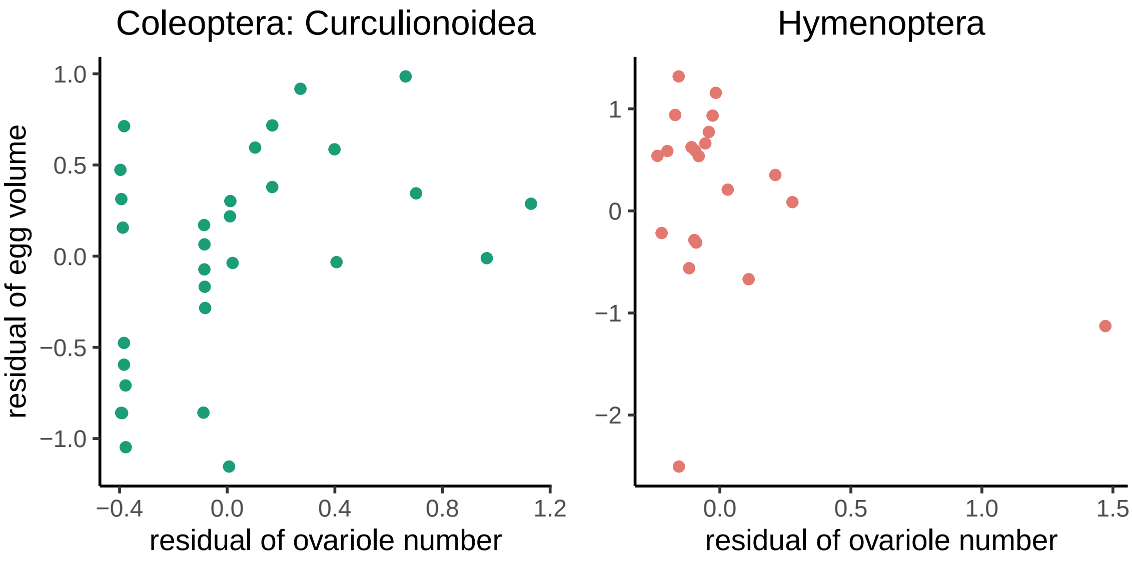

Figure S5: **Additional lineage-specific comparisons of egg volume vs ovariole number, phylogenetic residuals to body size, matching records at the genus level.** Weevils (Curculionoidea, left) were measured using elytra length and wasps (Hymenoptera, right) were measured using mesosoma width.

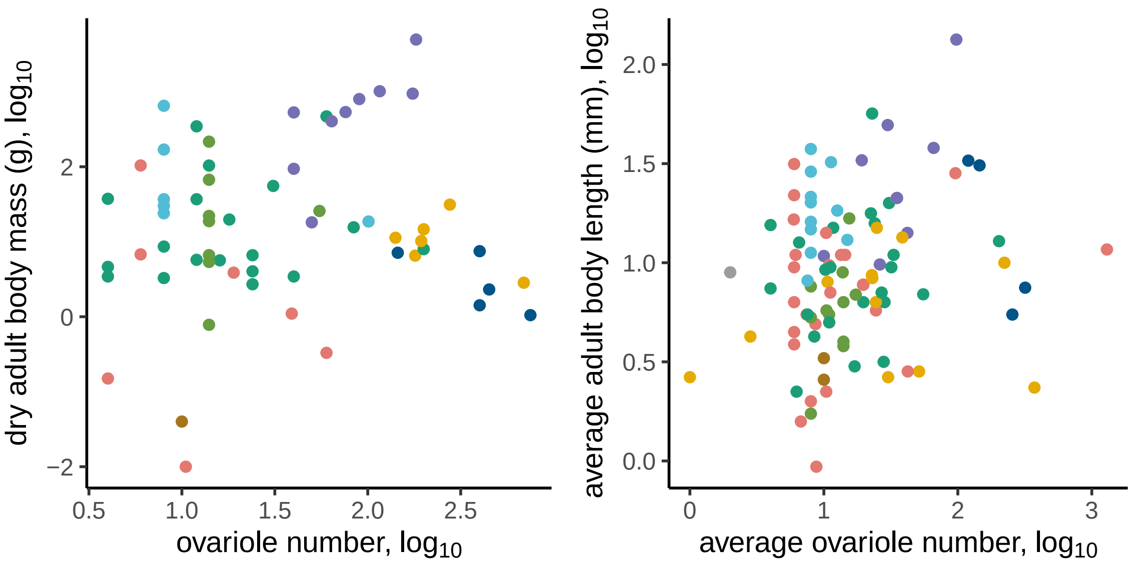

Figure S6: **Adult body size vs ovariole number.** Adult body mass compared at the genus level (left), and average adult body length compared at the family level (right).

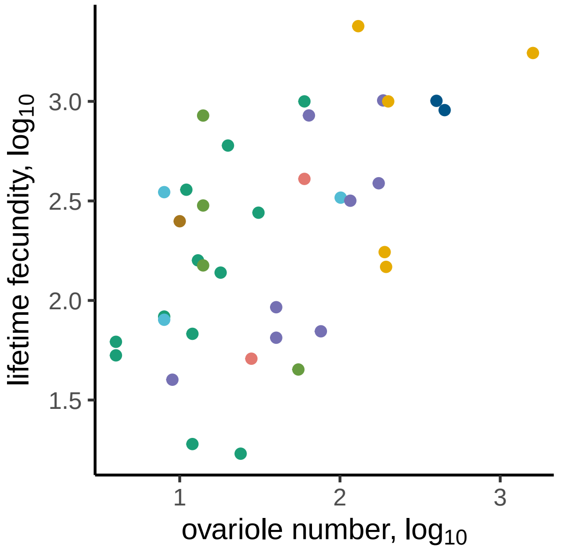

Figure S7: **Lifetime fecundity vs ovariole number, matching records at the species level.**

Table S1: Results of PGLS analysis of ovariole number and egg size across a posterior distribution

| analysis | taxonomic level | slope | MCC p-value | num. sig. / 1000 | taxa |
|---|---|---|---|---|---|
| ovariole number vs egg volume | species | -0.426 – -0.082 | 0.195 | 43 | 306 |
| ovariole number vs egg volume | genus | -0.356 – -0.128 | 0.066 | 470 | 482 |
| ovariole number vs egg volume, residuals to body mass | species | -0.646 – -0.333 | 0.003 | 833 | 24 |
| ovariole number vs egg volume, residuals to body mass | genus | -0.769 – -0.284 | 0.003 | 995 | 61 |
| ovariole number vs egg volume, residuals to body length | family | -0.685 – -0.304 | <0.001 | 966 | 98 |

Table S2: Results of PGLS analysis of ovariole number and egg size across a posterior distribution

| analysis | taxonomic level | slope | MCC p-value | num. sig. / 1000 | taxa |
|---|---|---|---|---|---|
| Drosophilidae ovariole number vs egg volume, residuals to thorax length | species | -0.814 – -0.799 | <0.001 | 1000 | 30 |
| Orthoptera ovariole number vs egg volume, residuals to body length | genus | -0.315 – 0.379 | 0.485 | 0 | 40 |
| Curculionoidea ovariole number vs egg volume, residual to elytra length | genus | -0.293 – 0.633 | 0.384 | 0 | 30 |
| Hymenoptera ovariole number vs egg volume, residuals to mesosoma width | genus | -2.131 – -0.288 | 0.139 | 13 | 21 |

Table S3: Results of PGLS analysis of ovariole number and body size across a posterior distribution

| analysis | taxonomic level | slope | MCC p-value | num. sig. / 1000 | taxa |
|---|---|---|---|---|---|
| ovariole number vs body length | species | 0.025 – 0.208 | 0.618 | 0 | 24 |
| ovariole number vs body mass | genus | 0.095 – 0.299 | 0.546 | 0 | 61 |
| ovariole number vs body mass | family | 0.123 – 0.177 | 0.031 | 29 | 98 |
| Drosophilidae ovariole number vs thorax length | species | 0.223 – 0.223 | 0.031 | 0 | 30 |
| Orthoptera ovariole number vs body length | genus | 0.132 – 0.450 | 0.001 | 993 | 40 |
| Curculionoidea ovariole number vs elytra length | genus | -0.211 – 0.257 | 0.917 | 0 | 30 |
| Hymenoptera ovariole number vs mesosoma width | genus | -0.112 – 0.355 | 0.482 | 0 | 21 |

Table S4: Results of PGLS analysis of ovariole number and fecundity across a posterior distribution

| analysis | taxonomic level | slope | MCC p-value | num. sig. / 1000 | taxa |
|---|---|---|---|---|---|
| ovariole number vs lifetime fecundity | species | 0.324 – 0.542 | 0.011 | 311 | 37 |
| ovariole number vs lifetime fecundity | genus | -0.275 – 0.601 | 0.002 | 267 | 65 |

### 4 Evolution of nurse cells

#### 4.1 Combining datasets

We used the descriptions of the mode of oogenesis recorded by Büning[33]. This author catalogued the ovary morphology for 136 insect genera, categorizing them into four modes: those without nurse cells (panoistic), with nurse cells adjacent to each clonally related, developing oocyte (polytrophic meroistic), with all nurse cells located in the germarium (telotrophic meroistic), and a unique mode of oogenesis reported only in Strepsiptera (reduced polytrophic meroistic ovaries).

Of the 136 genera observed by Büning, 70 are represented in the phylogeny used here[4]. Another 36 come from families or orders that have representative genera in the phylogeny, and within which all observations have the same recorded mode of oogenesis, when more than one was recorded. Therefore, we used a substitute genus as the phylogenetic tip for these 36 groups, bringing the total overlap between dataset and phylogeny to 106 taxa.

#### 4.2 Reconstructing evolutionary shifts in oogenesis mode

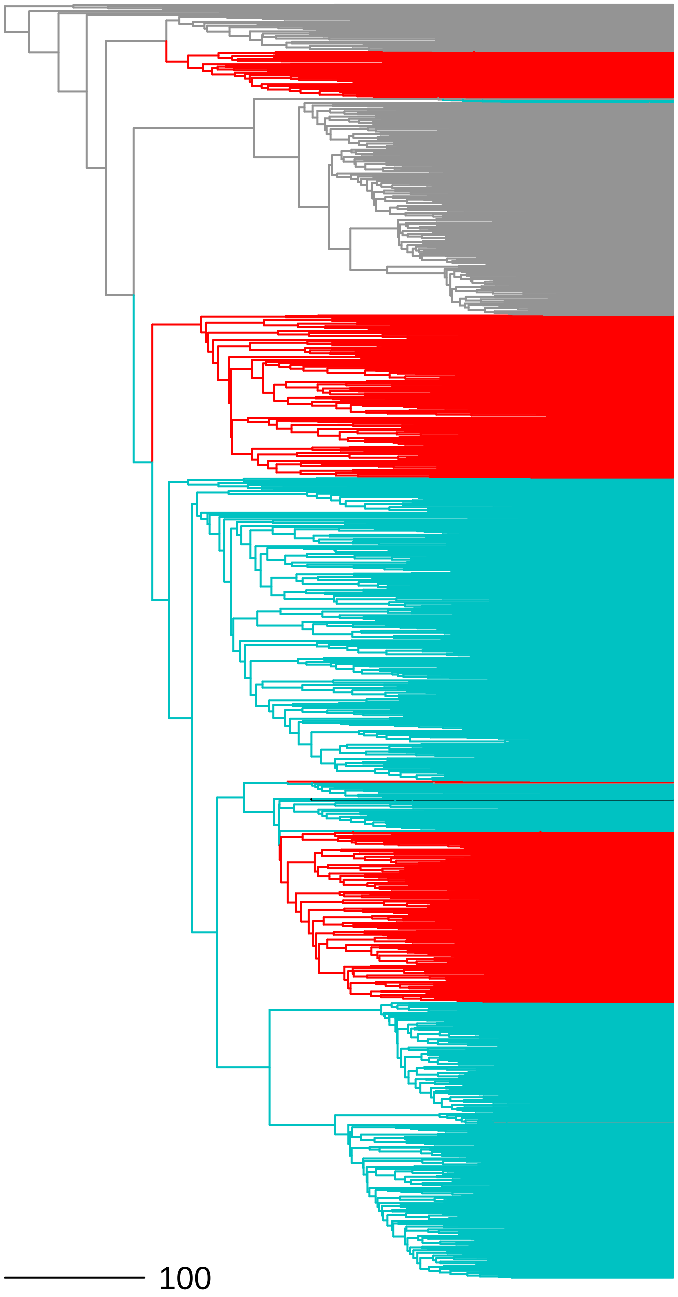

Figure S8: **Ancestral state reconstruction of oogenesis mode, full phylogeny.** Scale bar indicates 100 million years. Gray = panoistic, red = telotrophic meroistic, cyan = polytrophic meroistic, black = unique meroistic mode found in Strepsiptera.

Using these 106 records for ovary type, we reconstructed the ancestral state at each node of the published phylogeny of 1705 insect genera using an equal-rates model that allows for missing data (Fig. S8, R package corHMM, version 1.22[34] function rayDISC, node.states = 'marginal').

#### 4.3 Oogenesis mode model comparison

Table S5: Average corrected AIC (AICc) value from model comparison

| analysis name | BM1 | BMS | OU1 | OUM |
|---|---|---|---|---|
| ovariole number | 477.20195 | 473.03372 | 479.22592 | 482.42889 |
| egg volume | 3295.74949 | 3281.71001 | 3297.75729 | 3299.52446 |
| egg aspect ratio | -1486.86444 | -1506.89208 | -1484.85630 | -1483.01934 |
| egg asymmetry | -763.10658 | -785.44797 | -808.70150 | -810.07385 |
| egg curvature | 62.36224 | 35.70801 | 63.46489 | 64.20869 |

Table S6: Results of model comparison analysis over posterior distribution, showing the number of iterations out of 100 where the difference in model fit ($\Delta$AICc) was greater than 2.

| analysis name | BMS vs. BM1 | OU1 vs. BM1 | OUM vs. BM1 | OUM vs. OU1 | taxa |
|---|---|---|---|---|---|
| ovariole number | 90 | 0 | 0 | 0 | 506 |
| egg volume | 100 | 0 | 0 | 0 | 1567 |
| egg aspect ratio | 100 | 0 | 0 | 0 | 1488 |
| egg asymmetry | 100 | 100 | 100 | 20 | 844 |
| egg curvature | 100 | 1 | 1 | 1 | 781 |

Using the ancestral state reconstruction of evolutionary shifts in the mode of oogenesis, we inferred the most likely mode of oogenesis for all nodes and unobserved extant tips in the phylogeny (Fig. S8). We then compared the fit of models of trait evolution that take into account these shifts in oogenesis mode against those that do not. These comparisons were performed with the R package OUwie (version 1.57)[35].

Each analysis compared four models of evolution: single-rate Brownian Motion (BM1), multi-rate Brownian Motion (BMS), single-optimum Ornstein-Uhlenbeck (OU1), and an Ornstein-Uhlenbeck model with different optima for each mode of oogenesis (OUM).

These comparisons were repeated 100 times over a posterior distribution of trees. At each iteration we selected a random representative trait record for each genus in the phylogeny, when multiple records were available.

### 5 Modeling rate of ovariole number change

#### 5.1 Parametric bootstrap of Brownian Motion model

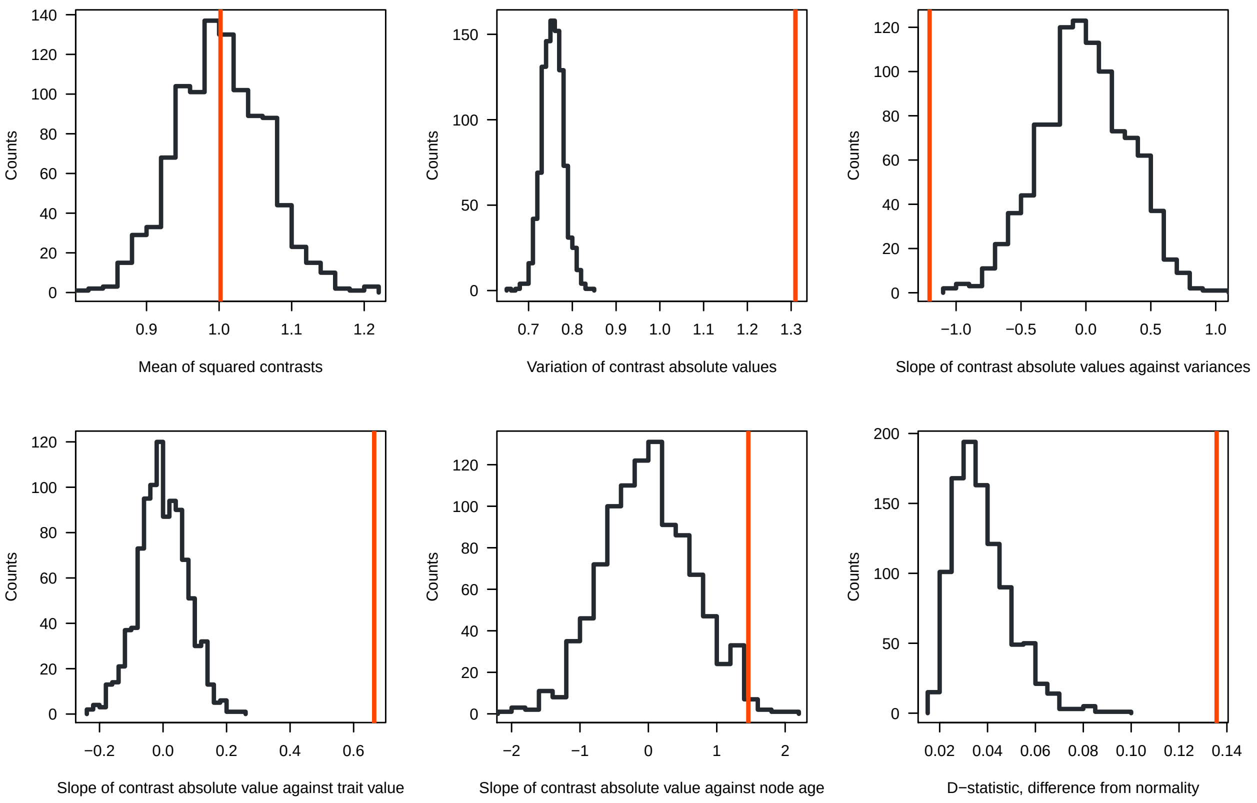

Figure S9: **Bootstrap analysis of Brownian Motion model for ovariole number evolution, using the R package arbutus.** In each panel the red line represents the observed value and the black distribution represents the bootstrap simulation. See Section 5 for details on each parameter.

We evaluated the fit of a Brownian Motion (BM) model for ovariole number evolution using the R package arbutus (version 0.1[36], Fig. S9). In this approach, a BM model is fit to the data (R package geiger, version 2.0.7)[31], and the resulting parameters of the model are used to simulate 1000 new datasets. Six statistical parameters are used to compare the phylogenetic contrasts of the observed data to the simulated data, and their interpretations are as follows[36]:

1. *Mean of squared contrasts.* The rate of evolution of ovariole number can be well estimated by the Brownian Motion model (the observed value falls within the null distribution).
2. *Coefficient of variation of the absolute value of the contrasts.* There is substantially more variation in contrasts than expected by chance, indicating heterogeneity in the rate of evolution beyond what a single-rate Brownian Motion model predicts (the observed value falls well outside the null distribution).
3. *Slope of a linear model fitted to the absolute value of the contrasts against their expected variances.* Contrasts are larger than expected on short branches in the phylogenetic tree, resulting in a negative slope. This could be explained by error in estimation of branch lengths.
4. *Slope of a linear model fitted to the absolute value of the contrasts against the ancestral state at the corresponding node.* The number of ovarioles is more correlated with contrast values than would be expected by chance. Phylogenetic nodes with a low ovariole number experience lower rates of evolution.

5. *Slope of a linear model fitted to the absolute value of the contrasts against node depth.* Contrast values are not correlated with time, falling within the null distribution. Therefore the rate of ovariole number change is not increasing or decreasing over time.
6. *The D statistic from a Kolmogorov-Smirnov test comparing the distribution of contrasts to an expected normal distribution.* The data do not fit a normal distribution of contrasts well, suggesting there are likely non-Brownian motion based processes at play (e.g. jump-diffusion processes).

#### 5.2 Assessing rate heterogeneity

Given the result that our dataset contains substantial rate heterogeneity, we identified regions of the tree with high and low rates of ovariole number evolution using the software BAMM (version 2.5.0)[37]. For this analysis, we calculated the average ovariole number for each genus in the insect phylogeny[4]. Average ovariole number was $\log_{10}$ transformed, and the tree was filtered to include only tips for which there were corresponding ovariole number data (sample size = 508). We used the R package BAMMtools (version 2.1.7)[38] to select priors, and ran BAMM for the maximum number of generations ($2 * 10^9$), sampling every $10^6$ generations.

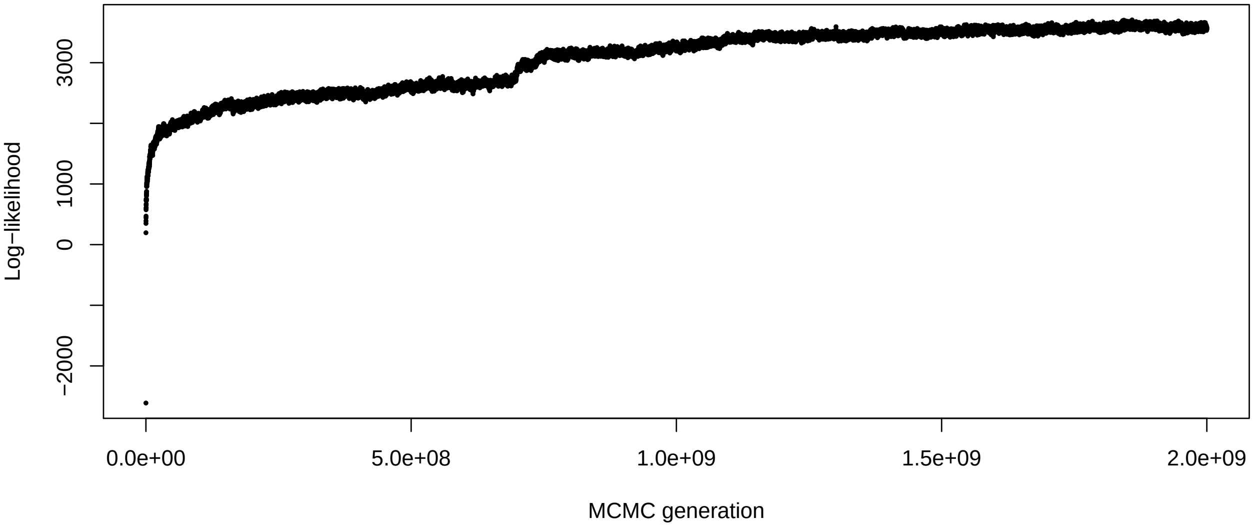

Figure S10: Convergence of trait diversification rate analysis

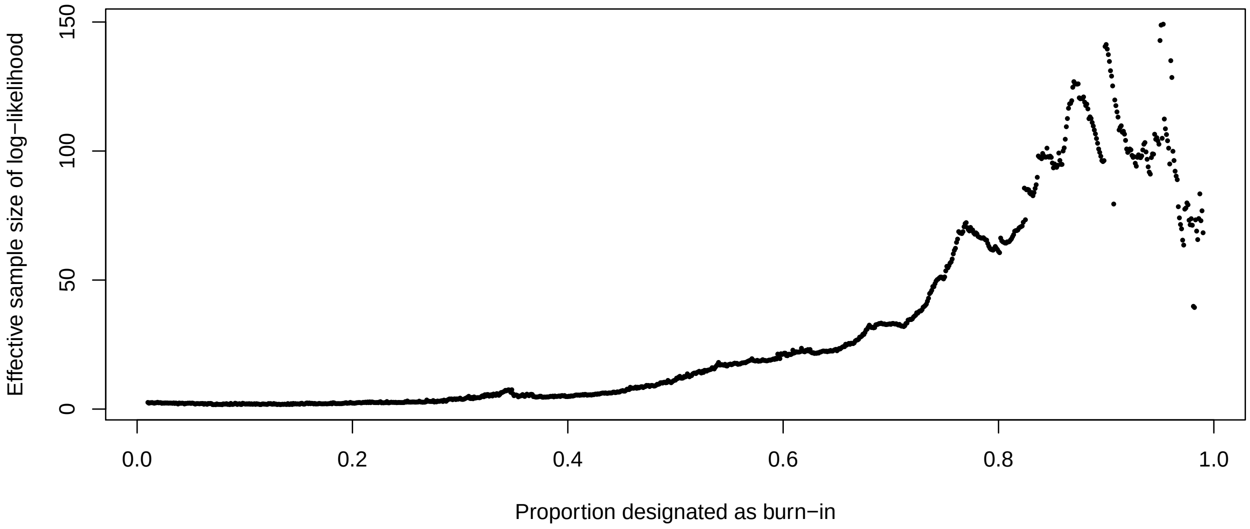

Figure S11: **Comparison of burn-in proportions.** The burn-in proportion that maximized the effective size was used in subsequent analyses.

Convergence was evaluated both visually (Fig. S10) and numerically by comparing the effective sample size for number of shifts and log-likelihood to the standard recommended by the software (>200). We determined the most appropriate burn-in proportion to use by finding the maximum effective sample size of the log-likelihood across an array of possible burn-in proportions (Fig. S11). Running BAMM for the maximum possible number of generations and selecting the optimum burn-in (Fig. S11) resulted in an effective size for the number of shifts of 482.51, and for log-likelihood of 149.15. Repeated BAMM analyses showed similar distributions of high and low rate regimes, indicating the implications for ovariole number evolution are robust to uncertainty in rate estimates. See Supplemental Methods Section 5.2 for details.

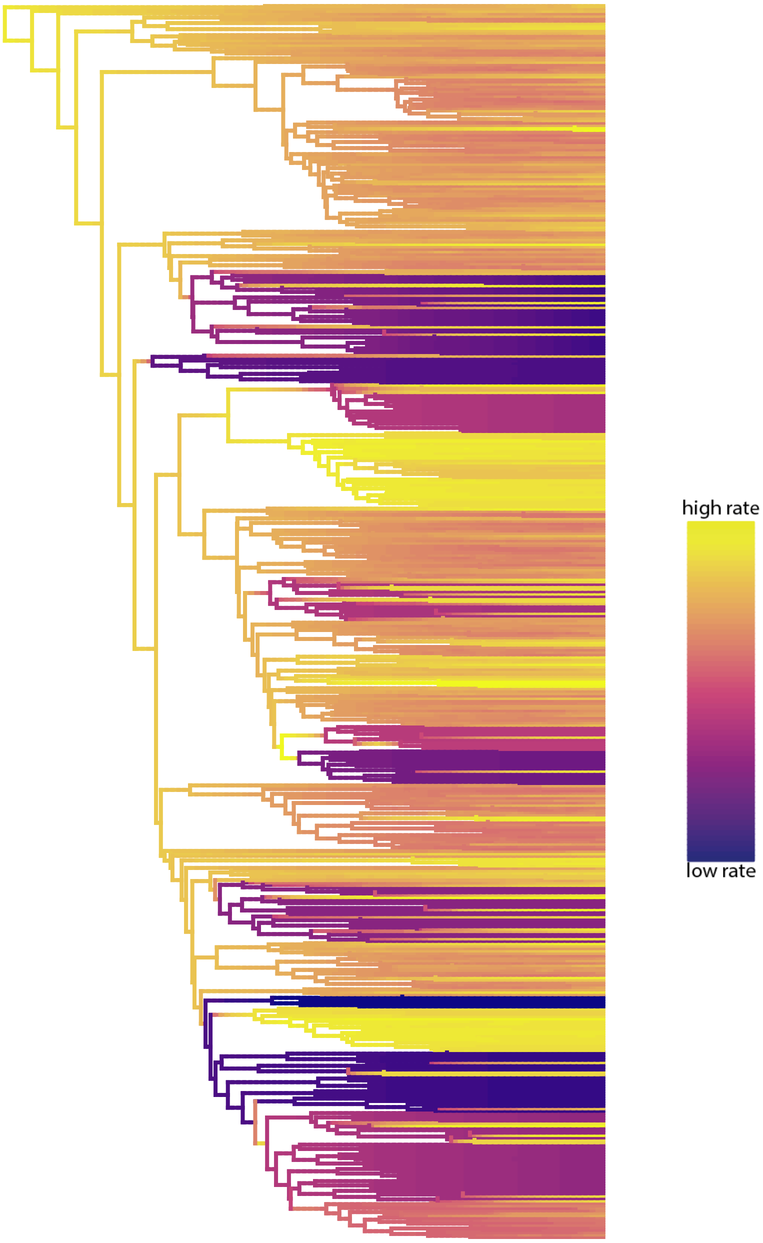

Figure S12: **Best rate shift configuration from BAMM trait diversification analysis on ovariole number.** Purple = low rate of evolution, yellow = high rate of evolution.

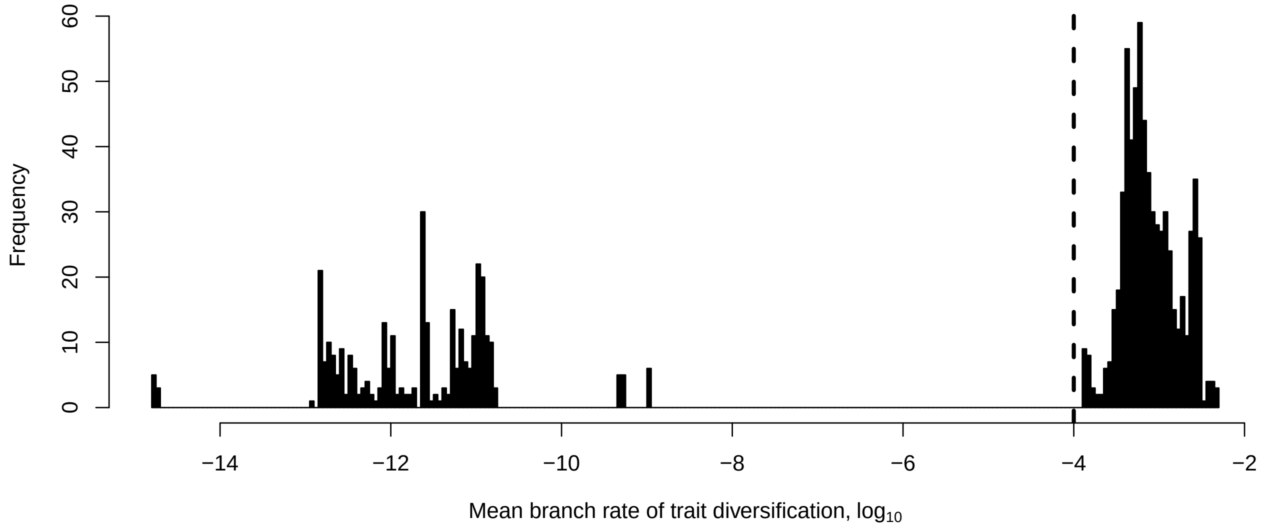

Figure S13: **Distribution of trait diversification rates.** Dotted line shows threshold used to assigned rate regimes.

The best configuration of rate shift regimes shows multiple independent clades with very low rates of evolution (Fig. S12). Visualizing the distribution of mean rates along branches revealed a discontinuous distribution, with one peak at a moderate rate of evolution and several clusters at extremely low rates, separated from the first peak by over six orders of magnitude (Fig. S13). We used this visualization to establish a threshold ($10^{-4}$) for assigning a binary rate regime to each node in the phylogeny, categorizing them as above (variable) or below (invariant) a threshold that separates these two peaks.

#### 5.3 Rate model comparison

We tested whether a BM model of evolution that incorporates the binary state (variable or invariant) as independent rate regimes can better explain the distribution of ovariole numbers than a single rate BM model, by comparing model fit using the R package OUwie (version 2.5)[35]. We find that a multi-rate model is significantly favored over a single-rate model ($\Delta$ AICc 1770.93).

#### 5.4 Comparing rates of trait diversification

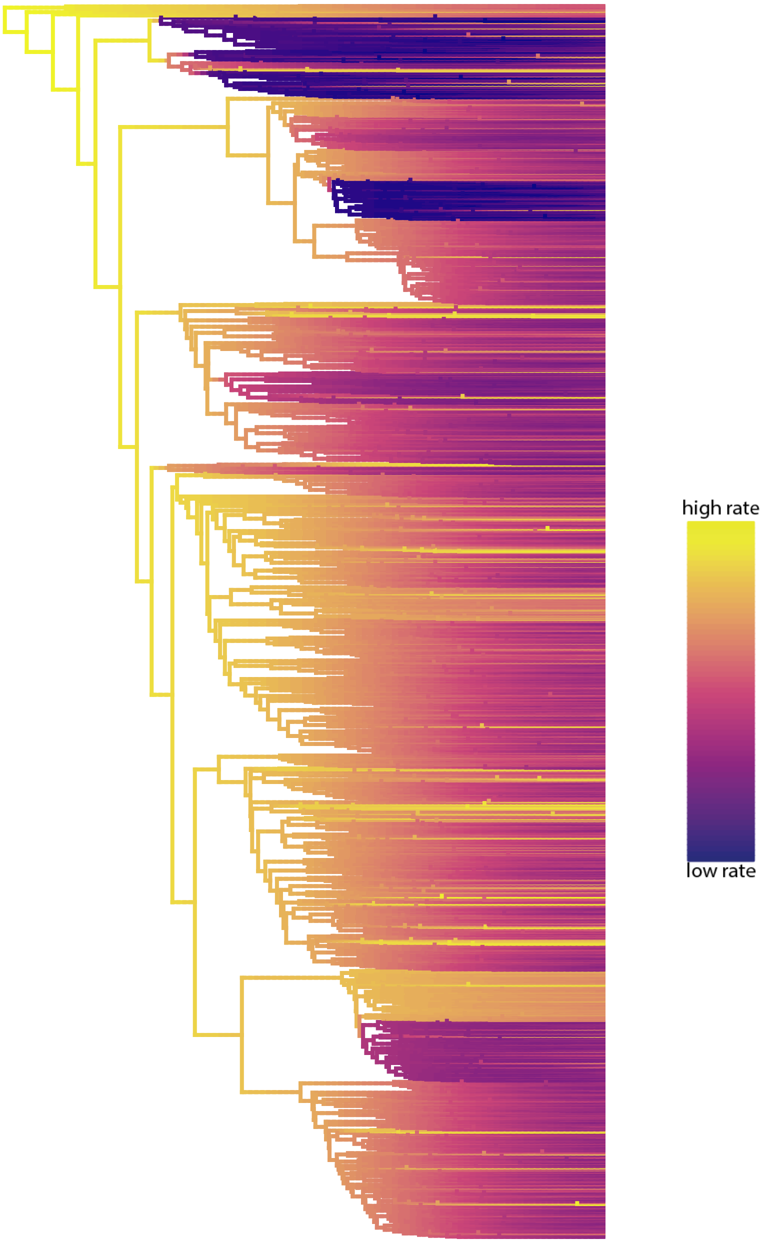

Figure S14: **Best rate shift configuration from BAMM trait diversification analysis on egg volume.** Purple = low rate of evolution, yellow = high rate of evolution.

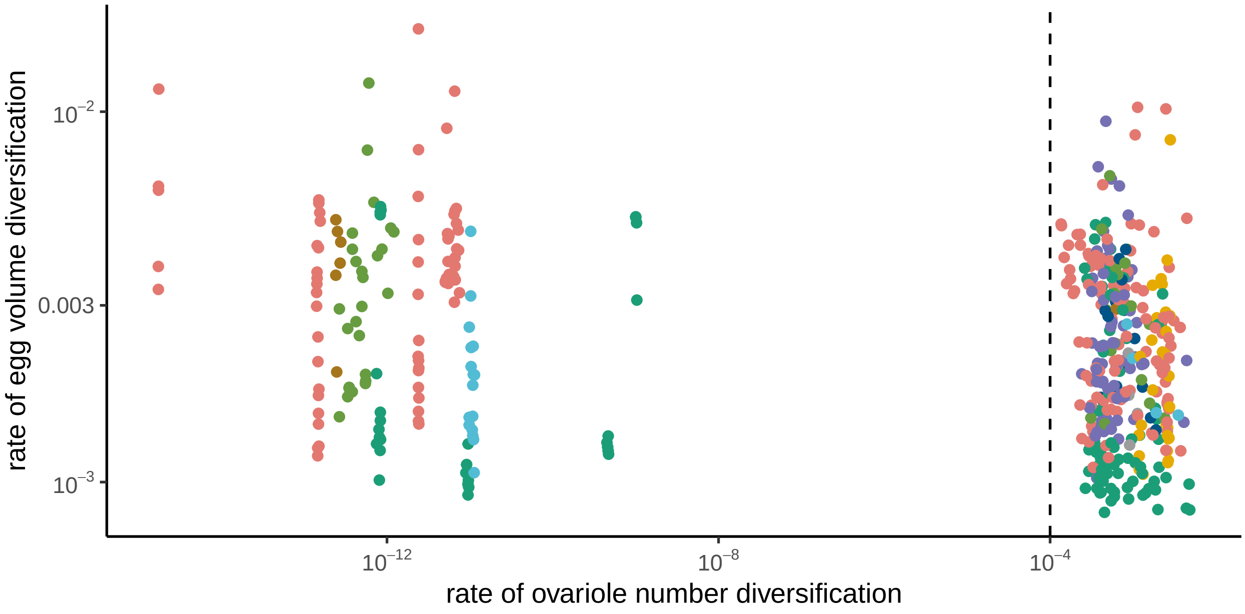

Figure S15: **Rates of trait diversification of egg volume and ovariole number.** Points are colored by phylogenetic groups shown in Fig 1a. Dotted line shows threshold used to assigned rate regimes.

We assessed the rate of egg volume diversification across insects using the same method of assessing rate heterogeneity as described in Section 5.2 (Fig. S14). This analysis converged in $7 * 10^9$ generations (the effective size for the number of shifts and log-likelihood were >200, 670.2 and 753.42 respectively). We compared the correlation between the rates of trait diversification for ovariole number and egg volume by matching the mean rate predicted for each insect genus (the tips of the phylogeny from the BAMM analyses, Fig. S15).
